# But the clock, tick-tock: the preeminence of relaxed clock models in total-evidence dated phylogenetics

**DOI:** 10.1101/2025.02.28.640870

**Authors:** Nicolás Mongiardino Koch, Jeffrey R. Thompson, Rich Mooi, Greg W. Rouse

**Affiliations:** Scripps Institution of Oceanography, University of California San Diego, La Jolla, California, USA; Department of Biology, Colorado State University, Fort Collins, Colorado, USA; School of Biological Sciences, University of Southampton, Southampton, UK; School of Ocean and Earth Sciences, University of Southampton, Southampton, UK; California Academy of Sciences, San Francisco, California, USA

**Keywords:** Echinoidea, total evidence, tip dating, relaxed clocks, fossilized birth-death, time calibration

## Abstract

Phylogenetic clock models translate inferred amounts of evolutionary change (calculated from either genotypes or phenotypes) into estimates of elapsed time, providing a mechanism for time scaling phylogenetic trees. Relaxed clock models, which accommodate variation in evolutionary rates across branches, are one of the main components of Bayesian dating, yet their consequences for total-evidence phylogenetics have not been thoroughly explored. Here, we combine morphological, molecular (both transcriptomic and Sanger-sequenced), and stratigraphic datasets for all major lineages of echinoids (sea urchins, heart urchins, sand dollars). We then perform total-evidence dated inference under the fossilized birth-death prior, varying two analytical conditions: the choice between autocorrelated and uncorrelated relaxed clocks, which enforce (or not) evolutionary rate inheritance; and the ability to recover ancestor-descendant relationships. Our results show that the latter has no impact on either topology or node ages and highlight a previously unnoticed interaction between the tree and clock models, with analyses implementing an autocorrelated clock precluding the recovery of direct ancestry. On the other hand, tree topology, fossil placement, divergence times, and downstream macroevolutionary inferences (e.g., ancestral state reconstructions) in sea urchins are all strongly affected by the type of relaxed clock implemented. In regions of the tree where molecular rate variation is pervasive and morphological signal relatively uninformative, fossil tips seem to play little to no role in informing divergence times, and instead passively move in and out of clades depending on the ages imposed upon them by molecular data. Our results highlight the extent to which the phylogenetic and macroevolutionary conclusions of total-evidence dated analyses are contingent on the choice of relaxed clock model, highlighting the need for either careful methodological validation or a thorough assessment of sensitivity. Our efforts continue to illuminate the echinoid tree of life, supporting the erection of the order-level clade Apatopygoida to include three living species last sharing a common ancestor with other extant lineages in the Jurassic. Furthermore, they also illustrate how the phylogenetic placement of extinct clades hinges upon the modelling of molecular data, evidencing the extent to which the fossil record remains subservient to phylogenomics.

---

A central tenet of evolutionary biology is that closely related species are more similar to each other than distantly related ones, an observation widely accepted by early naturalists whose generative mechanism was formalized by Darwin in the form of his theory of descent with modification (Darwin 1859). With the progression of time, similarities inherited from common ancestors become eroded as lineages experience independent evolutionary processes; those surviving constitute the data used to infer their phylogenetic relationships (Cracraft and Donoghue 2004; Maddison et al. 2007). Although some degree of proportionality between time and amount of change was rapidly accepted, its converse, i.e., the use of observed differences to date divergences, was only first proposed by Zuckerkandl and Pauling (1962). The “molecular evolutionary clock” (Zuckerkandl and Pauling 1965), as it later became known, posited that the constancy of substitution rates among lineages and through time, coupled with paleontological calibrations, could be used to infer divergence times from amounts of change in molecular sequences. One of the direct results of this development was the ability to impart a geologic timescale to phylogenetic trees (Morgan 1998).

Early attempts to derive evolutionary timescales were limited in numerous ways: they made use of a fixed topology, resulting in overconfidence of divergence times due to a lack of propagation of topological uncertainty; they enforced a strict clock, making the unrealistic assumption of a total absence of rate variation; and took the paleontological record at face value, directly applying fossil ages to internal nodes when these can only provide partial information to date cladogenetic events. These limitations have since been superseded by methods of simultaneous Bayesian inference that rely on relaxed clocks and implement prior calibration densities (Sanderson 1997; Rambaut and Bromham 1998; Thorne et al. 1998; Yoder and Yang 2000; Benton and Ayala 2003; Graur and Martin 2004; Hedges and Kumar 2004; Drummond et al. 2006; Yang and Rannala 2006; Ho and Phillips 2009).

While absent from their original formulation, molecular clocks were soon interpreted as an expected outcome of the predominant neutrality of molecular evolution (Kimura 1969). From this perspective, there seemed to be no room for morphological traits—whose evolution is primarily adaptive (Rieseberg et al. 2002; Ho et al. 2017)—to contribute to the inference of evolutionary timescales, even though such potential had been originally recognized (Mayr 1963; Simpson 1965). However, molecular clocks soon lost their preeminence in the neutralist-selectionist debates and became recognized instead as an emergent property of evolving biological systems, one whose usefulness remained even if it behaved only as an approximate, and often non-neutral, clock (Zuckerkandl 1987; Bromham and Penny 2003; Kumar 2005; Ho 2020). Soon after the adoption of statistical models of morphological evolution (Lewis 2001; Lee and Worthy 2012; Wright and Hillis 2014), researchers began inferring time-scaled trees (or chronograms) using morphological clocks, either on their own (Beck and Lee 2014; Lee et al. 2014) or alongside molecular ones (Pyron 2011; Ronquist et al. 2012a; Lee 2016).

The ability to infer chronograms that include extinct terminals revolutionized the fields of systematics, paleobiology, and macroevolution (Lee and Palci 2015; Hunt and Slater 2016; López-Antoñanzas et al. 2022). Node calibrations can only incorporate a small fraction of the fossil record’s temporal information, while a precise phylogenetic placement for fossils that are fragmentary or display intermediate morphologies can be hard to ascertain—caveats that are alleviated by the joint inference of topology and node ages using dated tips (i.e., ‘tip dating’; Ronquist et al. 2012; O’Reilly et al. 2015). The development of the fossilized birth-death (FBD) process (Heath et al. 2014), which models the history of diversification underlying the tree structure, further provided a natural and mechanistic framework to combine extant and fossil taxa and also allowed for the inference of ancestor-descendant relationships (Pett and Heath 2020; Wright et al. 2022). As genetic data became widely available, the inference of chronograms using a combination of molecular, morphological, and stratigraphic information, and sampling across both living and extinct lineages—known as total-evidence dating (Zhang et al. 2016)— became the gold standard for the inference of phylogenetic, biogeographic, and macroevolutionary history (Ronquist et al. 2012a, 2016; Arcila et al. 2015; Lee 2016; Gavryushkina et al. 2017; Pyron 2017; Lee and Yates 2018; Simões et al. 2020b; May et al. 2021; Mongiardino Koch and Thompson 2021; Troyer et al. 2022; Faurby et al. 2024; Tejada et al. 2024).

Total-evidence dating (TED) relies on a tripartite modelling framework (Warnock and Wright 2020) that includes a model of diversification (or tree prior, typically an FBD process), a model of trait evolution, and a (usually relaxed) clock model. Recent years have seen extensive efforts devoted to test the first of these (Matschiner 2019; Luo et al. 2020; O’Reilly and Donoghue 2020; May et al. 2021; Barido-Sottani et al. 2023; Simões et al. 2023b; Zhang et al. 2023), as well as to characterize the behavior of models of morphological change (Wright et al. 2016; Klopfstein et al. 2019; Rosa et al. 2019; Khakurel et al. 2024; Mulvey et al. 2024; Wright and Wynd 2024). The impact of implementing alternative relaxed clock models within a TED framework has received comparatively less attention. For example, the choice between autocorrelated and uncorrelated clock models, which differ on whether they assume or not, respectively, the existence of evolutionary rate inheritance, can dramatically impact timescales inferred by node dating molecular trees (Lartillot et al. 2016; Reis et al. 2018; Mongiardino Koch and Milla Carmona 2024). How they affect key aspects of TED inference such as fossil placement, recovery of ancestral-descendant relationships, and rates of morphological change, remains relatively unexplored.

Echinoderms represent an ideal empirical system for phylogenetically grounded studies of macroevolution and paleobiology (e.g., Hopkins and Smith 2015; Wright 2017; Lam et al. 2021; Mongiardino Koch 2021a; Wright et al. 2021; Thuy et al. 2022). Their complex skeleton, composed of multiple elements of durable calcitic stereom (in some lineages forming tightly interlocked structures, such as the echinoid test and the crinoid calyx), endows them with both a highly informative morphology and a remarkable preservation potential. The resulting wealth of echinoderm paleontological data can be readily integrated with information for extant lineages, producing robust morphological phylogenies extending back to Mesozoic or Paleozoic times (Kroh and Smith 2010; Thuy and Stöhr 2016; Rahman et al. 2019; Souto et al. 2019; Fau and Villier 2023; Saulsbury and Baumiller 2023). Nonetheless, only one echinoderm TED analysis has been performed to date—the echinoid (sea urchin) phylogeny of Mongiardino Koch and Thompson (2021).

Our goal here is to revisit and build upon the aforementioned study, leveraging state-of-the-art methodologies with extensive morphological, molecular (both transcriptomic and Sanger-sequenced), and stratigraphic datasets to infer the phylogenetic relationships and divergence times of all major living and extinct lineages of Echinoidea. Furthermore, we explore the sensitivity of our results to the choice of different parametric conditions, including different relaxed clock models and tree priors, highlighting the extent to which TED phylogenies and downstream macroevolutionary conclusions are impacted by modelling assumptions.

## Materials & Methods

### A) Dataset Construction

#### i. Morphological data

The morphological matrix for this study is taken from the seminal work of Kroh and Smith (2010), which targeted all family-level clades of post-Paleozoic echinoids using the type genus of each for character scoring. These taxa are currently classified at various taxonomic ranks, so we redefined the operational taxonomic units (OTUs) of this dataset as the most inclusive echinoid clades in the World Register of Marine Species (WoRMS Editorial Board 2024) containing a single coded species each. For the most part, these corresponded to either families, subfamilies, or tribes, although a handful of OTUs represent genus-level clades. In the original study, six families were excluded due to their incomplete or aberrant morphology, while a further five were removed after analyses to improve resolution. Here, we reincorporate the latter in the hopes that their placement will be more stable in a TED framework. We further added *Lissodiadema lorioli* Mortensen, 1903 as a terminal given that Lissodiadematidae is currently recognized at the family level (as the sister group of Diadematidae; WoRMS Editorial Board 2024), which further prompted a revision of the scorings of *Diadema antillarum* (Philippi, 1845). We also removed *Poriocidaris purpurata* (Thomson, 1872), a taxon now placed within the genus *Histocidaris*, leaving the latter represented only by *H. elegans* (A. Agassiz, 1879). Following Hopkins and Smith (2015) and Mongiardino Koch and Thompson (2021), we removed three characters from the original matrix and edited five others to correct errors. The final matrix, available in the Supplementary Material, contained 169 OTUs and included the late stem-group Archaeocidaridae as the single morphological outgroup (Thompson et al. 2020). A total of 303 morphological characters were employed, describing traits variable and diagnostic at higher taxonomic levels, yet largely invariant within OTUs. As in Kroh and Smith (2010), sixteen characters were treated as ordered.

#### ii. Molecular data

We incorporated two molecular datasets: the phylogenomic matrix of Mongiardino Koch et al. (2022) and a newly-gathered dataset of Sanger-sequenced loci. The first was downsampled by retaining loci with an occupancy above 80%, and then sorted using *genesortR* (Mongiardino Koch 2021b) in the R statistical environment (R Core Team 2024). We retained the 25 loci exhibiting the highest phylogenetic usefulness, defined using a multivariate approach that simultaneously targets high phylogenetic signal and low evidence of systematic biases (Mongiardino Koch 2021b; Mongiardino Koch and Thompson 2021). Whenever multiple terminals were available for a given morphological OTU, the highest occupancy one was kept. The sea cucumber *Holothuria forskali* Delle Chiaje, 1824 was used as the single molecular outgroup. Empty positions in the alignments left behind after taxon deletion were removed. The final phylogenomic dataset included 5,017 amino acids, spanning 33 of the 77 extant echinoid OTUs (see Fig. 1).

**Figure 1:**
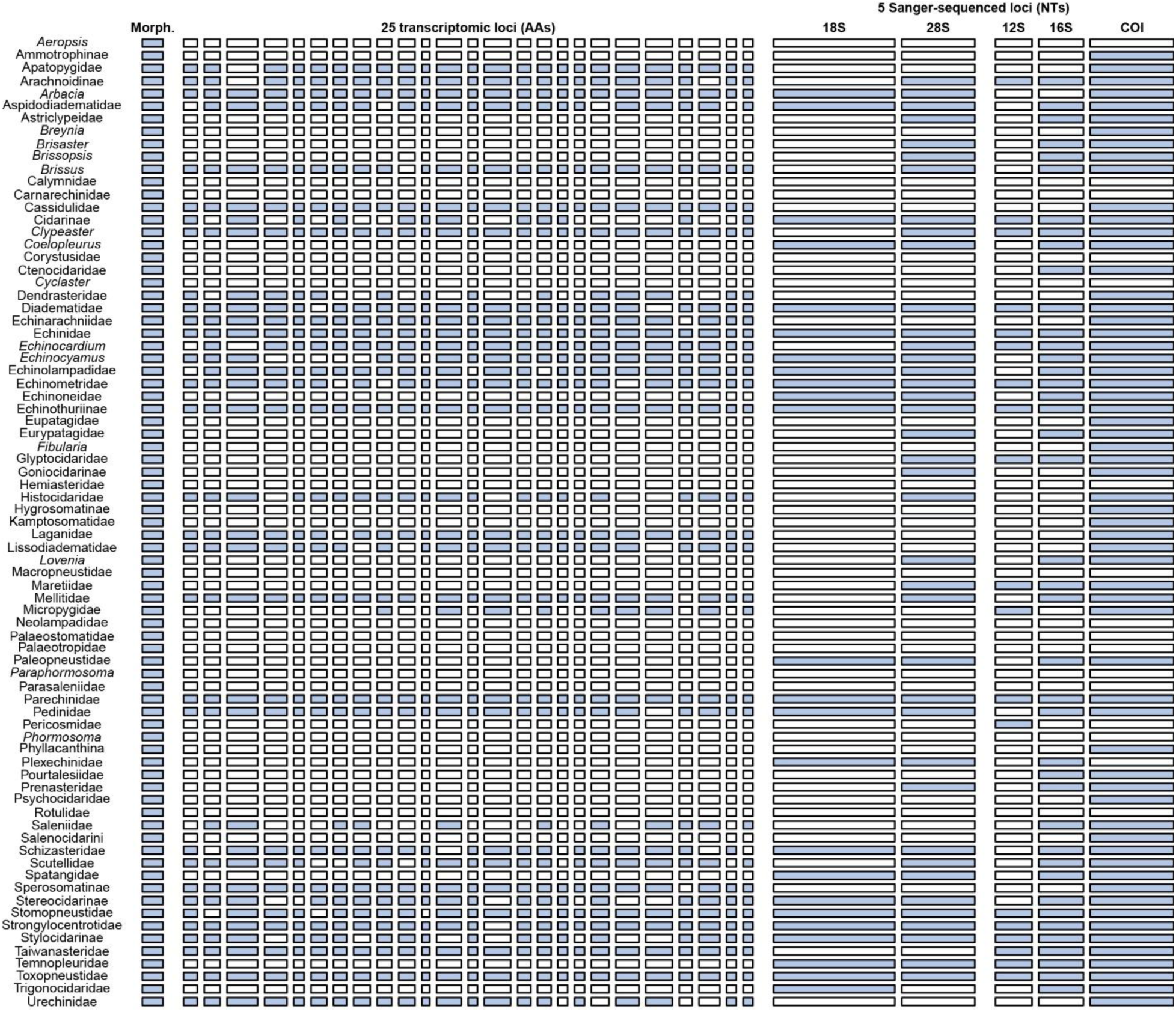
Total-evidence dataset of major echinoid clades assembled for this study. Columns represent the different matrices combined, including (from left to right) morphology, 25 transcriptomic loci coded as amino acids, and 5 Sanger-sequenced loci coded as nucleotides (nuclear: 18S rRNA and 28S rRNA, mitochondrial: 12S rRNA, 16S rRNA, and COI). Bar widths are scaled to the number of characters within each; blue bars represent partitions with data. Rows represent each of the 77 extant echinoid OTUs. The complete dataset further included the extant sea cucumber *Holothuria forskali* (coded only for molecular partitions) and 92 extinct echinoid clades (coded only for morphology). Final OTU names represent the least inclusive echinoid taxa encompassing all species from which data were gathered. Information about each sequence (species, GenBank/SRA accession number) can be found in Table S1.

To build a new dataset incorporating publicly available Sanger-sequenced data, all sequences of taxonomically non-redundant echinoids were downloaded from GenBank in September, 2021. This database was clustered into homologous loci using the approach described in Yang and Smith (2014), as implemented by Picciani et al. (2018). Briefly, we performed an all-by-all BLAST search (Camacho et al. 2009) to obtain pairwise similarity scores, excluded results with less than 50% sequence overlap, and clustered the remainder using MCL v.14-137 (Van Dongen 2000) with an inflation index of 1.0. The taxonomic breadth of the resulting clusters was explored by automatically assigning sequences to the higher-level OTUs of the morphological matrix. This relied on the R script *match_seq_to_taxonomy.R* (available on the Dryad repository), which uses as input a list of targeted OTUs and a taxonomic hierarchy. The latter was generated by web scraping the WoRMS repository using *deWoRMR* (https://github.com/mongiardino/deWoRMR), producing a table of all clade names to which each echinoid species is assigned (scraped in July, 2021). The fraction of OTUs represented within each cluster was tallied, and those containing data for less than 10% were discarded. Remaining clusters were further reduced to one sequence per OTU by retaining the species taxonomically closest to those included in the other datasets (i.e., morphological or phylogenomic, with priority given to the latter). If more than one sequence was deemed equally close taxonomically, the longest was retained. Data handling relied on R packages *ape* (Paradis and Schliep 2019), *phangorn* (Schliep 2011), and *tidyverse* (Wickham et al. 2019).

Nine clusters remained, including sequences of the nuclear 18S and 28S ribosomal RNA (rRNA; one and three clusters, respectively), mitochondrial 12S and 16S rRNA (one cluster each), cytochrome oxidase I (COI; two clusters), as well as a cluster containing complete mitogenomes. Preliminary alignments using MAFFT v.7.525 (Katoh and Standley 2013) revealed the multiple clusters of 28S and COI to be composed of one or more small fragments entirely nested within a larger one; thus, the corresponding clusters were merged and the longest sequence per OTU was selected. Existing COI data were complemented with newly generated data for 9 OTUs (highlighted in Table S1), representing major echinoid lineages which lacked publicly available data. COI was amplified using primers and PCR settings detailed in Arndt et al. (1996), products were purified with ExoSAP-IT (USB Corporation, OH, USA) and sequenced by Eurofins Genomics (Louisville, KY, USA). Forward and reverse sequences were *de novo* assembled on Geneious Prime 2023.2.2 (http://www.geneious.com) under default settings. Consensus reads were generated after removing base pairs of poor quality and manually resolving most ambiguities. One extra COI sequence was mined from the transcriptome of *Echinarachnius parma* using the top BLAST hit against the COI sequence of the closely related *Dendraster excentricus*. These sequences are deposited under GenBank accession numbers PQ619100-PQ619108.

Finally, mitochondrial loci (12S, 16S, and COI) were concatenated respecting the conserved echinoid gene order (Perseke et al. 2010), combined with complete mitogenomes, and aligned using the E-INS-i algorithm. Nuclear rRNA datasets were separately aligned with the L-INS-i algorithm. Gappy columns were deleted using trimAl v.1.5.0 (Capella-Gutiérrez et al. 2009), using a gap threshold of 0.24 for mitogenomes and 0.60 for nuclear rRNAs. Although initial efforts sought to use the entire mitogenome alignment, run times proved exceedingly long, so the three genes with high occupancy (12S, 16S, and COI) were excised and used instead. The five Sanger-sequenced loci obtained with this approach were incorporated into the final dataset—represented in Figure 1—as 5,229 nucleotide positions. All molecular data employed is identified in Table S1.

#### iii. Stratigraphic data

OTUs with any amount of molecular data were considered extant, even when the type species used for morphological coding was a fossil. For the remaining 95 OTUs, fossil ages were obtained from the primary literature and used as tip dates in the analysis. Additionally, 11 node dates were compiled from previous paleontological or phylogenetic studies (Smith et al. 2006; Thompson et al. 2017, 2019; Erkenbrack and Thompson 2019; Mongiardino Koch and Thompson 2021; Mongiardino Koch et al. 2022) and enforced for well-established monophyletic clades. Stratigraphic information can be found in the Dryad repository.

### B) Total-Evidence Dated Inference

#### i. Models of character evolution

Optimal modelling and partitioning of each data type (morphology, amino acid, and nucleotide) was determined using ModelFinder (Kalyaanamoorthy et al. 2017) as implemented in IQ-TREE v.2.1.3 (Minh et al. 2020). Partition merging was activated (Chernomor et al. 2016), and best-fit models were selected based on Bayesian Information Criterion (BIC) scores. Only models implemented in MrBayes v.3.2.7a (Ronquist et al. 2012b) were considered.

For morphological data, model space was circumscribed to the Mk_v_ model (Lewis 2001) with and without gamma distributed rate variation; partitioning schemes explored were: 1) unpartitioned, 2) by number of states, 3) by anatomical region, 4) similarity-based, and 5) homoplasy-based. Anatomical partitioning followed the regions defined by Kroh and Smith (2010). Similarity-based partitions were established with *EvoPhylo* (Simões et al. 2023a), using a Gower distance matrix (after randomly resolving polymorphisms) and selecting the optimal number of PAM (partitioning around medoids) clusters—i.e., the one maximizing the silhouette width. Homoplasy-based partitioning was performed following the approach of Rosa et al. (2019). A maximum parsimony tree under implied weighting (IW; Goloboff 1993) was inferred with TNT v.1.5 (Goloboff and Catalano 2016) with a concavity constant of 3 and used to derive character scores. Partitions were built by lumping characters with identical IW scores after rounding to the nearest decimal.

As with molecular data, IQ-TREE was allowed to improve model fit by merging the individual partitions of each original scheme, and the final model comparison was based on the fit of the best 9 resulting schemes (unpartitioned + 4 original schemes + 4 merged schemes). Across these, the tree topology was fixed to that obtained using the unpartitioned scheme, and 10 replicates of the entire process were performed. Optimal partitioning and modelling of the morphological matrix was attained with the merged homoplasy-based scheme, which attained a median BIC weight of 1.0. Further details of morphological model comparison are shown in Figure S1. Across-partition rate variation was activated and assigned a flat Dirichlet prior.

#### ii. Clock and tree models

The fossilized birth-death process was employed as tree prior (Heath et al. 2014; Zhang et al. 2016), using an exponential prior with a rate of 10 for the speciation probability, flat beta priors for extinction and fossilization probabilities, and fixing the fraction of sampled extant taxa to the known value. Ten nodes were constrained (see above) and calibrated using broad offset exponential distributions that were parameterized so as to leave 5% prior probability of origination at times even older than conservative soft maxima. The same approach was used for the tree age prior. A molecular backbone was enforced for extant taxa using 17 partial constraints, forcing the recovery of relationships found in genome-scale analyses (Mongiardino Koch and Thompson 2021; Mongiardino Koch et al. 2022); fossil affinities and remaining relationships among extant clades were inferred from the data. Uniform priors accounting for stratigraphic uncertainties and fossil lineage durations were used to calibrate all 95 fossil tips (Barido-Sottani et al. 2020). A diversified taxon sampling strategy was implemented for extant terminals (Höhna et al. 2011), and two sampling strategies were explored for fossils: activating and deactivating their potential recovery as sampled ancestors (Gavryushkina et al. 2014; henceforth referred as SA and noSA, respectively). The latter was attained by setting to zero the probability of move proposals that would place fossils as direct ancestors, as described in Simões et al. (2020a).

Two types of relaxed clock were explored: an autocorrelated lognormal model (TK02; Thorne and Kishino 2002), which assumes evolutionary rates are inherited across lineages following a Brownian motion; and an uncorrelated independent gamma model (IGR; Lepage et al. 2007), according to which the evolutionary rates of ancestors and descendants are uncoupled. Across both, a broad uniform prior was set for the base rate (normal distribution with mean of 0.001 and standard deviation of 0.01), an exponential prior with rate of 10 was used for the clock variance, and a separate clock was implemented for each data type (morphological, amino acid, and nucleotide) by unlinking relative rates across the corresponding sets of partitions. While heterogeneous clocks might also exist within each character type (Duchêne et al. 2014; Lee 2016), we assumed this setting captured the most salient aspects of rate variation in our data.

#### iii. Phylogenetic runs

Four separate inference procedures were explored varying the type of relaxed clock (TK02, IGR) and the strategy for fossil sampling (SA, noSA) in MrBayes v.3.2.7a (Ronquist et al. 2012b). For each of these, six independent runs composed of four chains each were continued for either 100 million generations with a 25% burn-in fraction (IGR), or 150 million generations with a 50% burn-in fraction (TK02). Sampling was performed every 5,000^th^ generation, resulting in 10k posterior samples drawn from each run (60k for each of the four analytical conditions). Convergence and stationarity of the post burn-in phase was visually confirmed with Tracer v.1.7.1 (Rambaut et al. 2018) and *rwty* (Warren et al. 2017). The average standard deviation of split frequencies ranged between 0.008 and 0.011. Effective sample sizes (ESS) for all parameters were larger than 170 for TK02 runs and larger than 321 for IGR runs; ESS values for topologies were all above 500. Analyses were summarized using majority-rule consensus (MRC) and maximum clade credibility (MCC) topologies with median heights; the latter was obtained with TreeAnnotator v.10.5.0 (Drummond and Rambaut 2007) using a random subsample of 15,000 posterior trees per analysis (2,500 per run). Depending on the computational burden of downstream analyses, these were run using a set of posterior trees further downsampled to just 1,500 per phylogenetic analysis (e.g., treespaces and chronospaces, see below).

### C) Analyses of Tree Topology and Divergence Times

#### i. Treespaces

The topological variation induced by the two analytical settings that were changed (i.e., the type of relaxed clock and the strategy for fossil sampling) was explored using treespaces (Hillis et al. 2005; Smith 2022). The number of matching quartets for all pairs of posterior trees were counted and used to derive a symmetric quartet distance using package *Quartet* (Sand et al. 2014; Smith 2019). Distances were then projected onto a two-dimensional space using multidimensional scaling (MDS).

#### ii. Chronospaces

While treespaces can efficiently summarize topological variation, they cannot portray temporal variation. Efficiently summarizing node age variation is complicated within the context of an analysis that simultaneously estimates topology and divergence times, as posterior chronograms do not share the same set of nodes. We explored two possible solutions to this issue: A) using the entire tree sample and focusing only on the subset of nodes shared by all, and B) defining a subset of nodes whose frequency (i.e., posterior probability) exceeds a given threshold, and downsampling the trees to those simultaneously consistent with them all. Clade frequencies were tallied, and those not present in all trees (solution A), or exhibiting frequencies below either 0.95 or 0.90 (solutions B_1_ and B_2_, respectively) were collapsed. Trees were then downsampled to those retaining the maximum number of nodes (i.e., those consistent with all nodes above the chosen threshold), resulting in datasets of node ages composed of 6k trees/41 nodes (solution A), 3.52k trees/78 nodes (solution B_1_), and 1.74k trees/90 nodes (solution B_2_). To maintain consistency with the methodology implemented to build treespaces, pair-wise Euclidean distances were estimated for each dataset and decomposed into two dimensions using MDS. The space obtained for solution A was rotated and scaled using Procrustes superimposition to minimize the sum of squared distances between the coordinates in the chronospace and those in the treespace. These approaches relied on functions from packages *chronospace* (Mongiardino Koch and Milla Carmona 2024) and *vegan* (Oksanen et al. 2024).

#### iii. Other tree properties

Across all subsampled posterior trees, we counted the number of taxa placed as direct ancestors (i.e., those with terminal branches of length zero), as well as the overall posterior probability (pp) with which each fossil resolved as an ancestor. We also estimated their stratigraphic congruence using the modified Manhattan stratigraphic measure (MSM*; Pol and Norell 2001), as well as their tree balance (using Colless’ index), level of treeness (proportion of total tree length made up of internal branches), and average node support. All five properties were compared across analytical conditions using either analyses of variance (ANOVAs) or, in the case of the number of sampled ancestors, a generalized linear model with a Poisson error distribution. Code relied on functions from packages *strap* (Bell and Lloyd 2015) and *apTreeshape* (Bortolussi et al. 2006).

### D) Macroevolutionary Inferences

To assess the impact that alternative parameterizations have on downstream analyses, we assessed clade origination times, evolutionary rates, and patterns of morphological change for trees obtained under alternative clock models. For the first, we characterized the general tempo of echinoid diversification through the lens of lineage-through-time (LTT) plots (Harvey et al. 1994), as well as explored the inferred divergence times of some focal clades. For the second, we extracted branch-specific rates of evolution for each of the three data types and tested whether significant differences exist between regular and irregular echinoids—a pattern previously supported using morphological data (Hopkins and Smith 2015)—using Kruskal-Wallis rank sum tests.

Finally, we performed a series of ancestral state reconstructions (ASRs) on the MCC trees of the IGR-SA and TK02-SA analyses. We iterated through every morphological character and pruned the MCC trees to the subset of terminals represented by non-missing, non-polymorphic codings. If the resulting trees contained more than 30 taxa and included members of both Euechinoidea and Cidaroidea (the two sister clades that comprise crown group Echinoidea), model-averaged marginal ASRs were performed for each MCC tree, combining both equal rates and different rates models, and implementing the prior distributions of FitzJohn et al. (2009) for the root node. We then compared the states assigned to the nodes corresponding to the three highest-level crown group clades (Echinoidea, and its two descendants: Cidaroidea and Euechinoidea) across both MCC trees, exploring the impact that the implemented clock model has on the reconstruction of morphological evolution across deep time.

## Results

Exploration of the posterior trees obtained for the different analyses performed revealed that major changes in both topology and node ages occur when implementing alternative relaxed clocks. Inference under either autocorrelated (TK02) or uncorrelated (IGR) clocks explored largely disjunct areas of both treespace and chronospace (Figure 2), indicating a strong and systematic effect of this decision on phylogenetic results. Posterior trees obtained when enforcing a TK02 clock proved less variable, suggesting a narrower region of maximum posterior probability. In stark contrast, preventing fossils from resolving as direct ancestors had no impact on the inference of phylogenetic relationships or divergence times, with topologies including (SA) or not including (noSA) sampled ancestors entirely overlapping each other in both treespace and chronospace. These conclusions are not modified by exploring a larger fraction of nodes ages (Figure S2).

**Figure 2:**
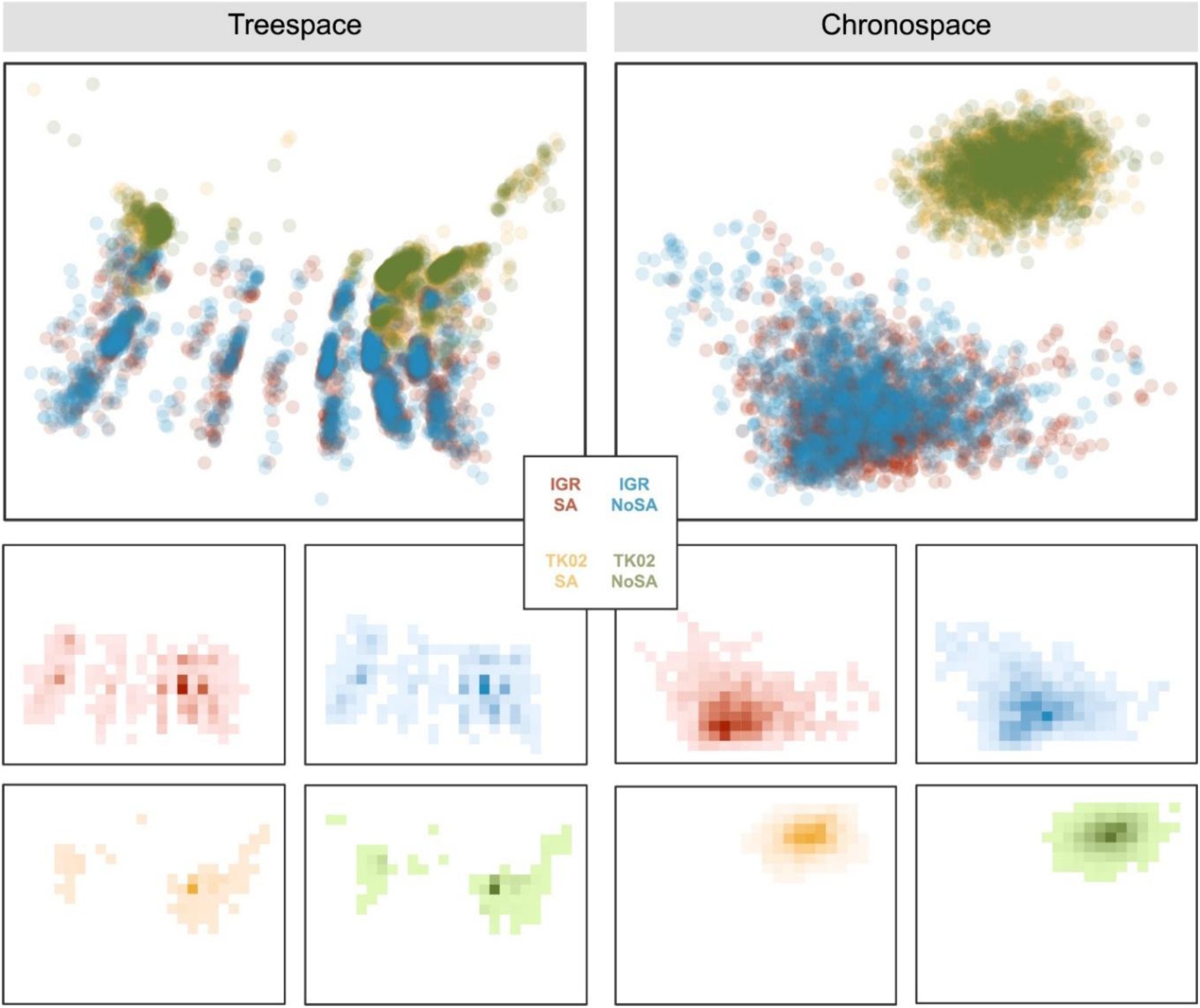
Treespace and chronospace condensing topological and temporal variability (respectively) across a random subsample of posterior trees obtained under the four analytical conditions explored. Analyses are colored as shown in the legend (center). The bottom panels show the same data using separate heatmaps for each analytical condition. While analyses implementing different clocks (TK02, IGR) resolve in different areas, those varying the fossil sampling strategy (SA, noSA) are completely superimposed, evidencing no effect of the latter on phylogenetic results.

Direct ancestry was relatively common among the posterior trees inferred using an uncorrelated relaxed clock (i.e., in the IGR-SA analysis). A median number of 9 terminals per posterior tree resolved as sampled ancestors of other clades, with a 95% confidence interval (CI) between 5 and 13 (Figure 3A). Nonetheless, as shown in Figure 2, IGR-SA and IGR-noSA analyses do not differ topologically or temporally; across these analyses, a fraction of fossil tips resolved as either direct ancestors or sister taxa of the same clades, without these alternative attachments to the same branch modifying topology or divergence times in any substantial way. Lineages involved in direct ancestor-descendant relationships under the IGR-SA analysis are summarized in Figure 4A. On the contrary, when an autocorrelated relaxed clock is enforced, no terminals are recovered as sampled ancestors, even when allowing for them (i.e., TK02-SA). In fact, the median posterior value for the parameter describing the proportion of fossils recovered as ancestors in this analysis is zero, compared to a median value of 0.095 (95% CI: 0.053-0.137) for the IGR-SA analysis. Given these results, subsequent comparisons focus only on the IGR-SA and TK02-SA trees. Further systematic differences are evident between these two analyses (Figure 3A), with posterior trees obtained under an autocorrelated relaxed clock exhibiting significantly lower stratigraphic congruence, level of treeness, and tree balance, while also showing significantly higher node support values (consistent with the reduced variability seen in the treespace of Fig. 2).

**Figure 3:**
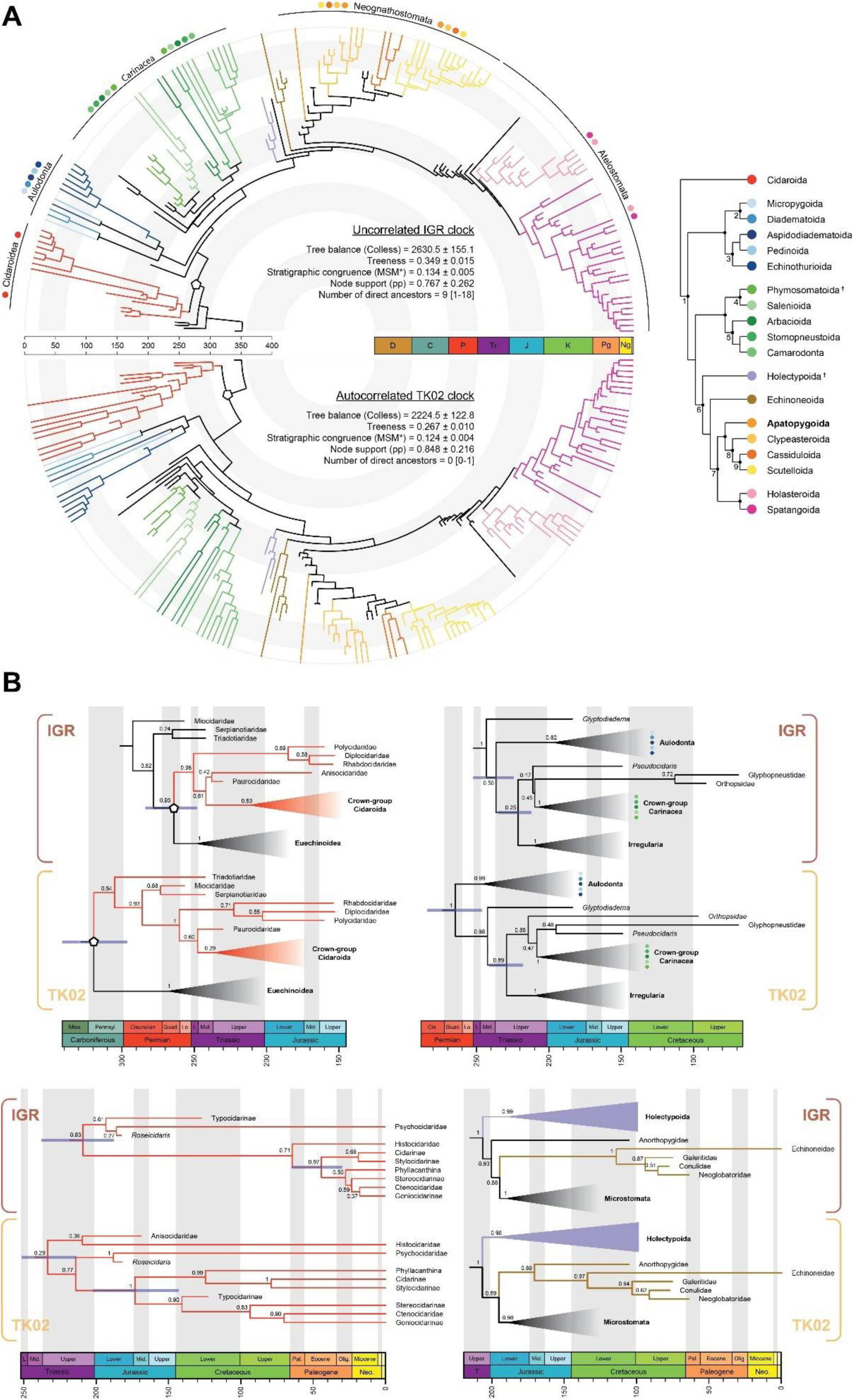
Comparison of TED chronograms obtained under uncorrelated (IGR-SA) and autocorrelated (TK02-SA) clock models. **A.** Summary of the differences between trees obtained under both analytical conditions. Chronograms depicted are MCC trees. Clades recognized at the level of order are colored and named on the cladogram (top right); other higher-level clades are shown using peripheral brackets or numbered nodes on the cladogram. Apatopygoida is shown in bold to highlight previously unrecognized status at the level of order. Properties estimated across both sets of posterior trees are shown in the center, as either mean ± standard deviation or median [range]. Here and throughout, a pentagon is used to identify the node of crown group Echinoidea. Numbered nodes on the cladogram: 1. Euechinoidea, 2. Diadematacea, 3. Echinothuriacea, 4. Calycina (sensu Kroh 2020), 5. Echinacea (sensu Kroh 2020), 6. Irregularia, 7. Microstomata, 8. Luminacea, 9. Echinolampadacea. Names for unnumbered nodes marked with a black circle are shown on the circular trees on the left. Crosses denote orders that are entirely extinct. **B.** Highlights of major differences between both MCC trees: top left, origin of crown group Echinoidea; top right, major euechinoid clades; bottom left, relationships and divergence times within crown group Cidaroida; bottom right, early diversification of irregular echinoids. Values on nodes are posterior probabilities; node age uncertainty for some clades of interest is shown using 95% highest posterior density intervals (see main text).

**Figure 4:**
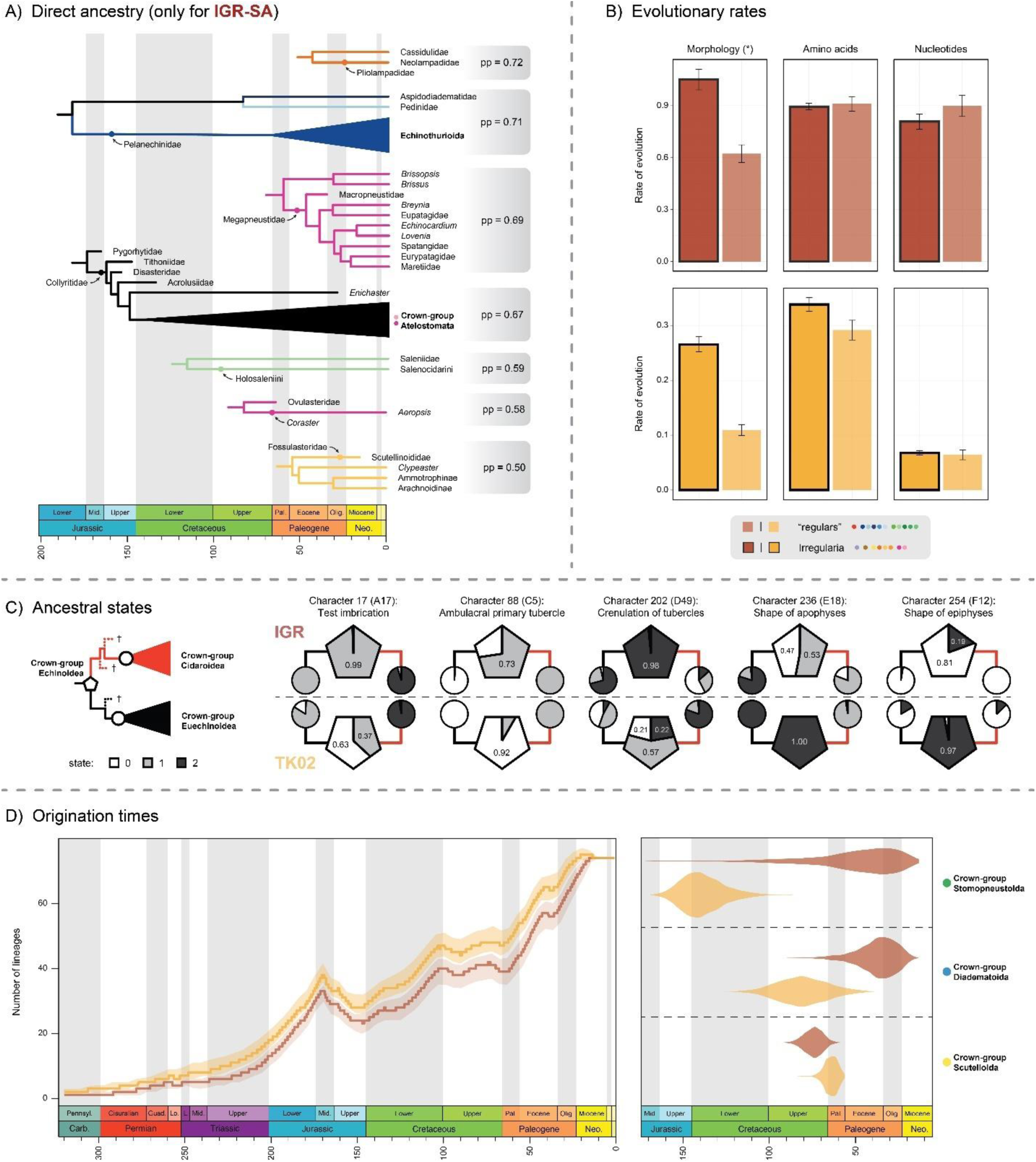
Macroevolutionary consequences of clock choice. **A.** Lineages with strong evidence (pp > 0.5) of being direct ancestors. While TK02-SA analyses allowed for ancestor-descendant relationships, none was recovered. Branch colors follow those of Fig. 3. **B.** Branch-specific evolutionary rates for the three data types (morphology, amino acids, and nucleotides), across “regular” echinoids and Irregularia. Significant differences (*P* < 0.05) are denoted by an asterisk. **C.** Ancestral state reconstruction of morphological traits for the nodes corresponding to the crown groups Echinoidea (pentagon), Cidaroidea (cirle, red branch), and Euechinoidea (circle, black branch), in the MCC trees of the IGR-SA and TK02-SA analyses. These nodes are identified as the last common ancestor shared by all extant descendants and might also include different numbers of fossil lineages (as shown in the diagram on the left using dotted branches leading to terminals marked with a cross). Values within pentagons represent maximum likelihood probabilities of each state. Characters are identified by their position in the morphological matrix used here, their code in Kroh and Smith (2010), and a brief description. Full details can be found in Fig. S5. **D.** Impact of the clock on reconstructed patterns of echinoid diversification, including a lineage-through-time plot (LTT, left), and posterior age densities for selected nodes (right). The LTT plot shows median node age values and LOESS-smoothed 95% confidence intervals. The posterior node age densities highlight three crown group orders which, depending on the clock implemented, are inferred to originate either before or after the Cretaceous-Paleogene (K-Pg) mass extinction event. Median inferred node ages for all comparable clades can be found in Figure S6.

Analyses further differed in crucial phylogenetic details, summarized in Figure 3B. Under an IGR clock, crown echinoids are dated to the Guadalupian (middle Permian), with a 95% highest posterior density (HPD) interval that encompasses post Permo-Triassic (P-T) mass extinction times. In contrast, a TK02 clock places this node decidedly within the Carboniferous, representing a difference in median inferred origination times of 55.6 Ma. Earlier times of origin are accompanied by the incorporation within the crown group of the late Permian/early Triassic members of the families Miocidaridae, Serpianotiaridae, and Triadotiaridae, lineages that otherwise resolve as stem group taxa (Fig. 3B, top left). TK02 analyses also result in older ages for the two main lineages of crown group echinoids—Euechinoidea and Cidaroidea—and within the latter, of Cidaridae, which is dated to either 172.2 Ma (95% HPD: 141.3-202.3 Ma; TK02) or 45.7 Ma (95% HPD: 30.7-65.8 Ma; IGR). These alternatives are also accompanied by changes in relationships, not just for fossil taxa (examples of which include the placement of *Glyptodiadema*, Anisocidaridae, and Typocidaridae) but also for extant lineages, including whether histocidarids or psychocidarids represent the sister group to the remainder of crown group Cidaroida. Within Irregularia, different clocks support the existence of either two clades of stem group irregulars—Holectypoida and Anorthopygidae—or just one, with anorthopygids resolving instead as an early member of the extant Echinoneoida (Fig. 3B, bottom right). Despite these striking differences, order-level relationships are consistent among analyses, and are summarized on the cladogram of Figure 3A. MCC and MRC trees for both IGR-SA and TK02-SA inference conditions, including 95% HPD intervals and posterior probabilities, can be found Figures S3 and S4, respectively.

A few other nodes that proved remarkably sensitive to the implemented clock are shown in Figure 4D, including three crown group clades that are dated to times at either side of the Cretaceous-Paleogene (K-Pg) mass extinction by different clocks. Although TK02 showed a general tendency to favor older divergence times, a handful of nodes are pushed forward under this clock, including that corresponding to crown group sand dollars (Scutelloida). Divergence times for all comparable clades across both clocks are shown in Figure S6. While the effect of the implemented clock model on branch durations is pervasive, it does not greatly modify general patterns of diversification (Fig. 4D) or estimates of evolutionary rate variation among major lineages (Fig. 4B). Irregular echinoids are confirmed to be characterized by significantly higher rates of morphological evolution (as also shown in Hopkins and Smith 2015), yet these are accomplished in the absence of elevated molecular rates. One downstream inference that did show high analytical sensitivity was that of the ancestral states reconstructed at the origin of crown group Echinoidea. Thirteen morphological traits differed in the most probable state assigned to said node, five of which are highlighted in Figures 4C (and fully explained in Figure S5). Not only do these trees imply different locations for the origin of key morphological innovations, but they occasionally also differ on which of the two major extant lineages bears the ancestral condition (for example, characters 88 and 236).

## Discussion

Bayesian methods of total-evidence dating that implement relaxed clocks and FBD processes are regarded as the gold standard for the simultaneous inference of tree topologies and divergence times, one that builds upon decades of methodological innovation. These approaches combine numerous benefits, including a reliance on mechanistic models describing all aspects of the underlying evolutionary dynamics, as well as an ability to combine all available sources of information (and uncertainty). They can also produce chronograms that sample thoroughly across both living and extinct diversity in a manner conducive to accurate macroevolutionary inferences. TED approaches have been thoroughly tested using simulations, as well as extensively used among vertebrates (e.g., Beck and Lee 2014; Arcila et al. 2015; Lee 2016; Pyron 2017; Gavryushkina et al. 2017; Lee and Yates 2018; Simões et al. 2020b; Tejada et al. 2024) and a handful of clades within arthropods (e.g., Ronquist et al. 2012a; Sharma and Giribet 2014; Zhang et al. 2016; Gustafson et al. 2017; Gainett et al. 2024) and plants (e.g., May et al. 2021; Coiro et al. 2023; Flores et al. 2023). Even though echinoderms offer unique advantages to utilize—and test—these methodologies, they remain relatively unexplored. Our goal here was to expand upon the only echinoderm TED analysis previously performed, the study of Mongiardino Koch and Thompson (2021) on Echinoidea, using improved methodologies, better sampled datasets, and an exploration of the effects of some analytical decisions.

The aforementioned study had incorporated transcriptomic data for 27.6% of extant terminals, a value that was increased here to 42.9%, and further raised to a value of 82.8% of extant terminals with some amount of molecular information through the addition of Sanger-sequenced data. This reveals the sustained relevance of these legacy datasets in the genomic era, especially for taxa that will remain difficult to sample and preserve in optimal conditions, as is the case for many deep-sea invertebrates. Mongiardino Koch and Thompson (2021) had also relied on a partitioning scheme for morphological data that was based on the number of character states (Gavryushkina et al. 2017; Mulvey et al. 2024), an approach that ensures that characters are assigned a transition matrix of appropriate dimensions (although see Khakurel et al. 2024), but can also suffer from issues of character weighting (King et al. 2017). Recent years have seen an appraisal of the importance of accurately modelling morphological data, as well as the development of new algorithmic approaches to character partitioning (Lanfear et al. 2017; Rosa et al. 2019; Azevedo et al. 2022; Casali et al. 2022; Simões et al. 2023a) that tend to fit the data better than either unpartitioned or anatomy-based partitioning. Our results broadly align with previous efforts, confirming that partitioning morphological data always improves model fit (Clarke and Middleton 2008; Tarasov and Genier 2015), and that homoplasy-based methods outperform other alternatives (see Fig. S1; Rosa et al. 2019; Casali et al. 2022; Lepeco and Melo 2024). The latter result makes sense, as the restriction of model space for morphological data to variants of the Mk renders partitioning an exercise in finding the best way of modelling among-character rate variation. Nonetheless, depending on the software employed, the need to ensure that transition matrices are correctly specified remains (MrBayes enforces this automatically). Finally, it should be pointed out that all original schemes were improved by a subsequent step of partition merging, which we argue should become standard as the modelling of morphological data continues to evolve to resemble that of molecular datasets, both in complexity and automatization.

The other main difference with respect to previous efforts was the implementation of an FBD tree prior, supplanting the use of the improper uniform tree prior (May et al. 2021). This prompted a number of decisions, among them the implemented strategy for fossil taxon sampling. FBD processes can infer ancestor-descendant relationships directly from the data, improving the modelling of diversification history for epidemiological and paleontological datasets for which sampled terminals do not necessarily arise through cladogenesis (Gavryushkina et al. 2014). Empirical efforts have shown that direct ancestors are likely common among known fossil species (Foote 1996), and that recognizing that speciation might occur through mechanisms other than strict bifurcation can affect our understanding of diversification dynamics (Crouch et al. 2021) and trait evolution (Caetano and Quental 2023). Nonetheless, direct ancestry should probably only be accommodated if one expects such relationship to be present in the data (Simões et al. 2020a), a probability that should decrease when sampling is sparse, focused on higher-level terminals, and spanning a vast period of time—conditions which are all met here. A similar logic applies to the choice of relaxed clock: autocorrelation (i.e., the inheritance of evolutionary rates) is a sensible biological expectation (Gillespie 1984; Tao et al. 2019), yet this pattern can be entirely eroded by a sparse taxon sampling. To our knowledge, only the morphological tip-dated study of sphenodontian reptiles by Simões et al. (2020a) had assessed sensitivity to both of these factors, and their concomitant effect on TED inference had never been explored before the present study.

Our results reveal a striking asymmetry in the impact exerted by these two decisions, with the type of clock greatly affecting both topology and divergence times, whereas accommodating direct ancestry shows no effect. Although the former has been long appreciated (e.g., Ronquist et al. 2012a; Zhang et al. 2016; Tejada et al. 2024), the latter is surprising. Previous analyses based on morphological datasets had found that allowing for direct ancestry substantially modified divergence times (Bapst et al. 2016; Simões et al. 2020a), resulting in older node ages as morphological change must be accommodated within fewer branches. While allowing for direct ancestry also had an effect on the age of one focal node in a TED analysis (Gavryushkina et al. 2017), the magnitude of change was comparatively small and in the opposite direction (i.e., favoring slightly younger nodes). It seems likely that as morphological data represents a progressively smaller fraction of total-evidence datasets, the alternative placement of a given fossil as either a direct ancestor or a sister clade will have less of an effect on the amount of change reconstructed for the attachment branch. While further research should clarify if this result is dataset-specific, we believe that—within the context of large-scale TED inferences—direct ancestry might be amenable to accommodation without affecting branch durations, and therefore have no consequence for divergence time estimation.

On the other hand, the recovery of ancestor-descendant relationships is entirely contingent on the relaxed clock implemented, revealing an extreme case of interdependence between the modelling components of Bayesian dating. Under the autocorrelated TK02 relaxed clock, only 0.1% of posterior trees contained sampled ancestors, and the few that did exhibited only a single ancestor-descendant pair; in comparison, all posterior trees contained sampled ancestors under the uncorrelated IGR clock, with numerous taxa placed with confidence as direct ancestors of other lineages (Figure 4A). It is worth noting that previous morphological phylogenies of major echinoid clades explicitly depicted both direct ancestry and budding cladogenesis (Smith 2007; Kroh and Smith 2010), the second of which could be reduced to direct ancestry if long-lived lineages were constrained temporally to the stratigraphic range of a single coded species, as done here. One such relationship implied by Kroh and Smith (2010) and corroborated here is that between Corasteridae and Aeropsidae (Figure 4A). Similarly, previous authors had suggested that Echinothuriidae arose “through” the Jurassic genus *Pelanechinus* (Durham and Melville 1957; Smith 2015), and highlighted the extent to which its morphology appeared to be “transitional” between lineages now recognized as the main branches of extant Aulodonta (Groom 1887; Philip 1965). These ideas are validated by our results, which place Pelanechinidae as a direct ancestor along the stem of Echinothurioida. Finally, the morphology of other terminals (e.g., *Fossulaster* and *Scutellinoides*) is similar enough that their recovery as ancestor-descendant pairs—given the fact that they were assigned disjunct stratigraphic ranges—constitutes a sensible outcome. Although a strict interpretation of direct ancestry might not always be warranted when working with terminals above the species level, the ability to infer trees in which lineages can explicitly evolve through sampled ancestral morphotaxa might still be beneficial, and is certainly consistent with how paleontologists have interpreted a number of these clades.

This ability, nonetheless, exists only if one assumes uncorrelated rates of evolution. A previous study on hymenopteran insects had noted that a much higher number of direct ancestors was obtained under an IGR clock compared to a TK02 clock (Zhang et al. 2016), although the value for the latter was not null, as it is here. This result was considered implausible and attributed to a high prior probability of direct ancestry under IGR that an uninformative morphological partition was unable to override. This explanation does not seem to apply here: the number of taxa involved in direct ancestry under IGR is reasonable, and our morphological dataset is highly informative, supporting topologies that are relatively consistent with phylogenomic ones (barring a handful of exceptions; Mongiardino Koch et al. 2018, 2022; Mongiardino Koch and Thompson 2021), and that exhibit levels of missing data among extinct taxa that are low and comparable to those of extant ones. Instead, we suggest an alternative explanation for this phenomenon: autocorrelated clocks might be unsuited for the inference of direct ancestry, as forcing evolutionary rates to change following a Brownian motion precludes the recovery of branches of zero length. Under this evolutionary scenario, a jump from a measurable (i.e., non-zero) ancestral effective branch length to a null one in a descendant (needed for the latter to be placed as a sampled ancestor), might be beyond what the model can allow. In this study, this would have to happen independently for all three implemented clocks (given that fossil terminal branches also have effective branch lengths for molecular partitions), something that the model does not seem to accommodate, precluding the inference of direct ancestry. The fact that this pattern had not been identified before might be partially due to the scarcity of TED analyses using amino acid data, which seem to exhibit significantly lower among-lineage rate variation than either morphology or nucleotides (Fig. S7), making large jumps in rates even less likely. Another contributing factor might be the lack of availability of some of these models across software. For example, most sampled-ancestor FBD analyses have been performed using the BEAST/BEAST2 family of software which have not consistently implemented autocorrelated clocks, thus preventing a thorough exploration of model sensitivity.

As already mentioned, the impact of the relaxed clock extends well beyond the recovery of direct ancestry, resulting in a different balance between stratigraphy and character data (King 2021), and having broad phylogenetic and macroevolutionary consequences (Figs. 2-4). We believe most of these to be driven by alternative clocks differing in the way they accommodate variation in molecular rates among lineages, not just because molecular data constitute most (97.4%) of the dataset, but because similar differences were found when using alternative clocks within the context of node dating (Mongiardino Koch et al. 2022). For example, median node ages for Cidaridae obtained here range between 44 and 173 Ma, values which are highly consistent with the age range of 51-147 Ma previously obtained for this clade when node dating under different clocks (Mongiardino Koch et al. 2022). The only factor in common between these studies is that inference was performed under both autocorrelated and uncorrelated clocks—Mongiardino Koch et al. (2022) did not incorporate fossils, had no morphological data, used a much larger molecular dataset, and employed both different software and tree prior. Cidarids (and potentially all cidaroids) have significantly lower rates of molecular evolution than all other extant echinoids (Mongiardino Koch et al. 2018). In the absence of node age constraints for the clade, their time of origin is mostly determined by the ability of the relaxed clock to successfully model shifts in evolutionary rates, a scenario under which uncorrelated clocks can favor dates that are systematically underestimated (Crisp et al. 2014). Although one would hope that the temporal information present in the tip ages of fossil cidaroids would help further constrain these dates, the general uninformativeness of cidaroid morphology (Hopkins and Smith 2015; Mongiardino Koch and Thompson 2021) causes these taxa to vary widely in their phylogenetic placement (Figure 3B), moving in and out of clades as these are dated to younger or older times. The same seems to happen with three fossil clades—Miocidaridae, Serpianotiaridae, and Triadotiaridae—variously considered by previous authors as either stem group echinoids, cidaroids, or euechinoids (and thus, among the earliest crown group lineages; Durham and Melville 1957; Philip 1965; Smith 1990, 2004; Kroh and Smith 2010; Mongiardino Koch and Thompson 2021). Their morphology is relatively consistent with any of these interpretations, so their position is mostly determined by the inferred ages of surrounding nodes. These various topological and temporal changes lead to alternative scenarios of the evolution of key aspects of echinoid morphology (Figs. 4C and S5). These include the nature of the perignathic girdle (the structure supporting the muscles used for feeding) in the most-recent common ancestor of extant echinoids, as well as whether or not their test was tightly-sutured (as that of most living forms), or relatively flexible and made of imbricating plates (as in many Paleozoic stem-group lineages). These different optimizations either validate or negate long-discussed putative synapomorphies of crown group Echinoidea, as well as inform on the extent to which living cidaroids, which have long been considered to display plesiomorphic character states, resemble the last common ancestor of crown group echinoids (Smith 1981, 1984; Smith and Hollingworth 1990; Thompson et al. 2015, 2020).

Despite all of these uncertainties, our results support a robust scheme of relationships among the major branches of the echinoid tree of life (Fig. 3A), as well as a consistent history of diversification through time (Fig. 4D). The recovery of the extant Apatopygidae as an early offshoot of irregular diversification is congruent with previous phylogenomic efforts (Mongiardino Koch et al. 2022), and confirms that the clade is unrelated to other extant “cassiduloids”, a result that likely was previously obtained due to the general difficulty of accurately placing living fossils (i.e., long-lived, slow evolving taxa) when performing morphological tip dating (Turner et al. 2017; King 2021; Mongiardino Koch and Thompson 2021; Mongiardino Koch et al. 2023). Although morphological similarities between apatopygids and nucleolitids had been highlighted as potential evidence of a close relationship (Mortensen 1948; Kier 1966; Suter 1994; Souto et al. 2019; Mongiardino Koch et al. 2022), these clades emerge as independent offshoots of the early neognathostomate diversification (as was also suggested by Kroh and Smith 2010). Accordingly, we erect the order-level clade Apatopygoida and follow Kroh and Mooi (2024) by assigning to it the family Apatopygidae, including its three extant species: *Apatopygus recens* (Milne Edwards, 1836), *Apatopygus occidentalis* H.L. Clark, 1938, and *Porterpygus kieri* (Baker, 1983). This taxonomic change also highlights the uniqueness of these species, which last shared a common ancestor with the remainder of extant Neognathostomata (sand dollars, sea biscuits, and allies) deep in the Mesozoic. As currently circumscribed, Apatopygidae also includes the Cretaceous genera *Jolyclypus* (two nominal species) and *Nucleopygus* (approximately 16 nominal species), as listed in Kroh and Mooi (2024).

While much work is still needed at lower taxonomic levels, few uncertainties remain regarding the emergence of the major clades of echinoids. Only one set of relationships has proved sensitive to the dataset and implemented methodology, that involving the earliest split among extant Carinacea. While Salenioida occupies that position here, in agreement with morphological signal (Kroh and Smith 2010), phylogenomic efforts revealed a stronger but ambiguous signal for Arbacioida to occupy that position (Mongiardino Koch et al. 2022). Furthermore, Echinoneoida remains the only order-level clade whose placement has not been addressed using genome-scale datasets, and for which more than one possible location has been previously obtained (as either the sister group to all other extant irregulars, or just to neognathostomates; Kroh and Smith 2010; Mongiardino Koch and Thompson 2021). While improvements in topological resolution among less divergent extant clades will advance as molecular data becomes more available (e.g., Lee et al. 2023; Minin et al. 2024), their interrelationships with extinct lineages will likely continue to hinge on modelling choices, especially for regions of the tree for which morphology has proven less informative.

## CONCLUSIONS

Here, we focused on combining large-scale morphological, molecular, and stratigraphic datasets for the major lineages of Echinoidea, and analyzing them using state-of-the-art phylogenetic methods. Despite working at the edge of what is currently computationally feasible within the realm of Bayesian total-evidence dating (involving year-long run times on a high-performance computer cluster), we find that key aspects of our results remain contingent upon methodological decisions that are seldom explored or justified. Within the context of tip dating, model selection approaches for validating the choice of relaxed clock have generally succeeded for morphological analyses (Lee et al. 2014; Dembo et al. 2016; Simões et al. 2020a); yet the few total-evidence studies that attempted this were met with uncertainty (Ronquist et al. 2012a). This highlights the need for a careful assessment of sensitivity; in fact, many of the results shown here to hinge upon the choice of clock model (e.g., the composition of the earliest splits within the echinoid crown group) were previously hailed by some of us as novel phylogenetic results made possible by total-evidence dating (Mongiardino Koch and Thompson 2021). The impact of this methodological choice goes beyond changes in tree topology and divergence times, as it can modify downstream macroevolutionary inferences, affecting our understanding of the congruence between phylogeny and stratigraphy, and even limiting our ability to infer ancestor-descendant relationships.

After molecular data grew in availability and genome-scale datasets assumed the role of basis upon which phylogenetic relationships, divergence times, and macroevolutionary patterns became routinely inferred, total-evidence dating was received as a means of integrating once again the irreplaceable information that the fossil record provides to understanding evolutionary history (Lee and Palci 2015; Pyron 2015). Our results challenge this view, showing instead that fossil terminals can sometimes remain largely relegated. This effect is evident for regions of the tree characterized by a relatively uninformative morphological signal coupled with stark variations in molecular rates. Under these conditions, characteristic of the echinoid stem group and early diversification of cidaroids, fossil tips fail to provide relevant information to constrain node ages, which become otherwise determined by the clock model. Our understanding of these key transitional fossils is thus entirely contingent upon assumptions regarding rate inheritance among characters they do not even possess.

## ACKNOWLEDGEMENTS

This manuscript benefitted from discussions about molecular clocks and direct ancestry with Tiago R. Simões and April M. Wright. We also want to thank Sonja Huč and Marina McCowin for DNA extractions and PCR amplifications, Natasha Picciani for help with the use of scripts for orthology inference, and Ferdinand Marlétaz and Yuwu Chen (San Diego Supercomputer Center) for computational assistance. The beginning of the title of this article is taken from the 1971 song ‘The Musical Box’ by Genesis.

## DATA AVAILABILITY

Newly generated molecular data is available from Genbank with accession numbers PQ619100-PQ619108. Accession numbers for all preexisting molecular data are listed on Table S1. The *deWoRMr* R script is available from GitHub: https://github.com/mongiardino/deWoRMR. All data and code used in this analysis are deposited on the Dryad Digital Repository: …

## Supplementary Material

**Figure S1:**
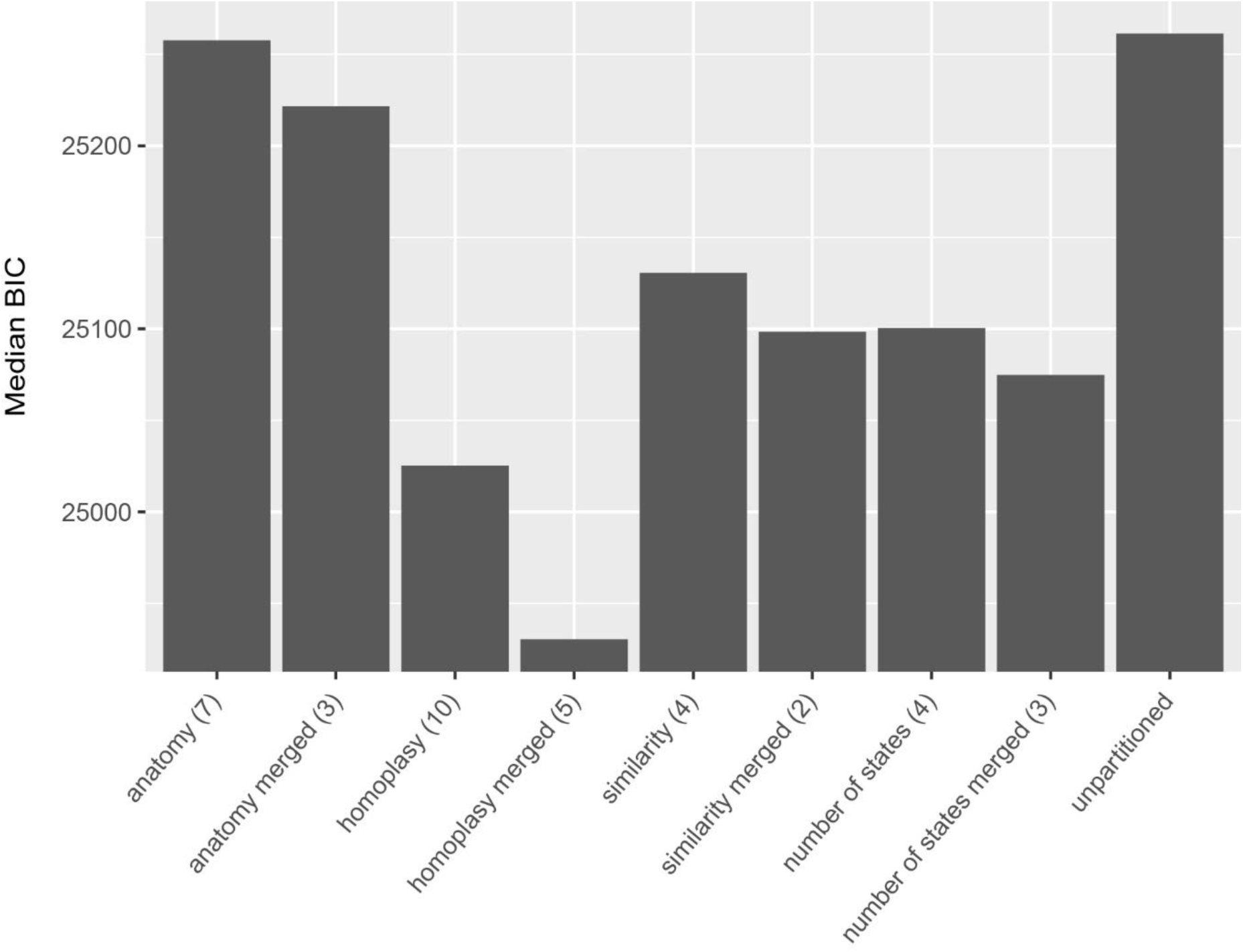
Fit (median BIC across 10 replicates) of alternative partitioning schemes for the morphological data. Each scheme shows the number of partitions in parentheses.

**Figure S2:**
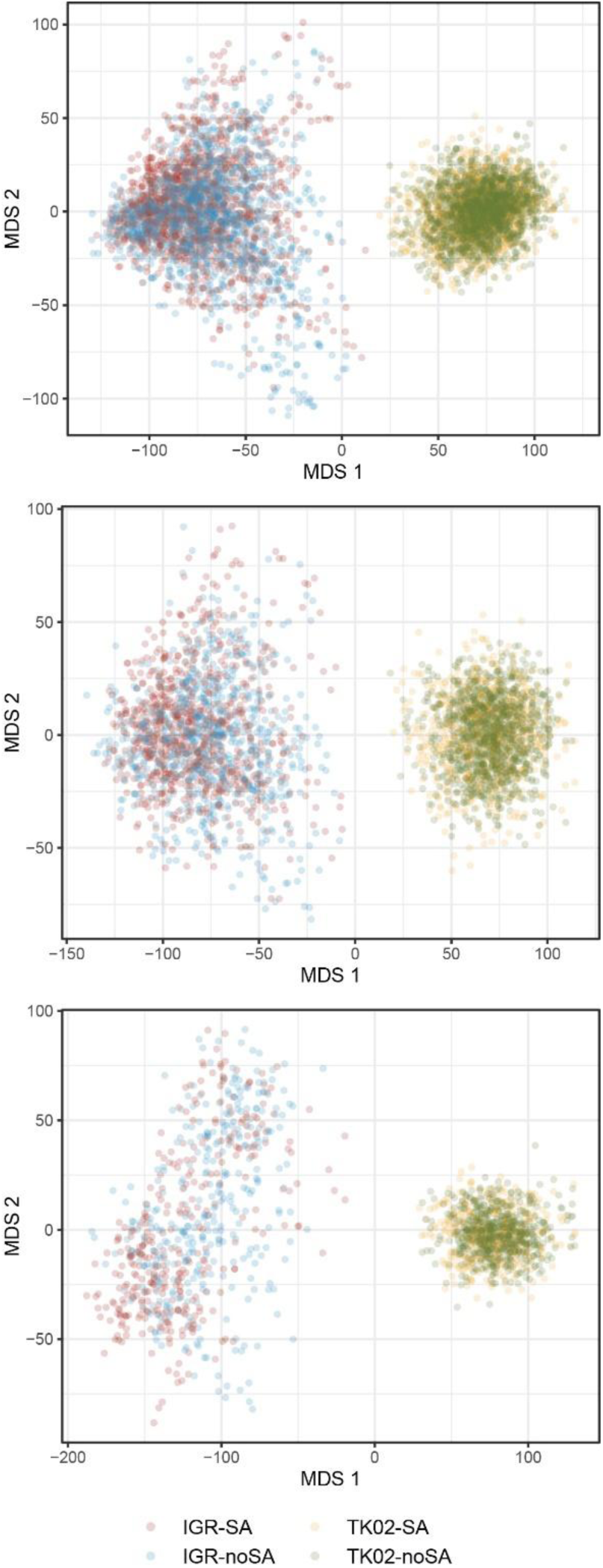
Chronospaces (two-dimensional multidimensional scaling of node ages for posterior trees) built using different numbers of nodes. **Top.** 41 nodes present across all sampled posterior trees. This is the same plot as shown in Figure 2, except that the original axes have not been rotated. **Middle.** 78 nodes with posterior probability > 0.95 (simultaneously present in 3.52k trees). **Bottom.** 90 nodes with posterior probability > 0.90 (simultaneously present in 1.74k trees).

**Figure S3:**
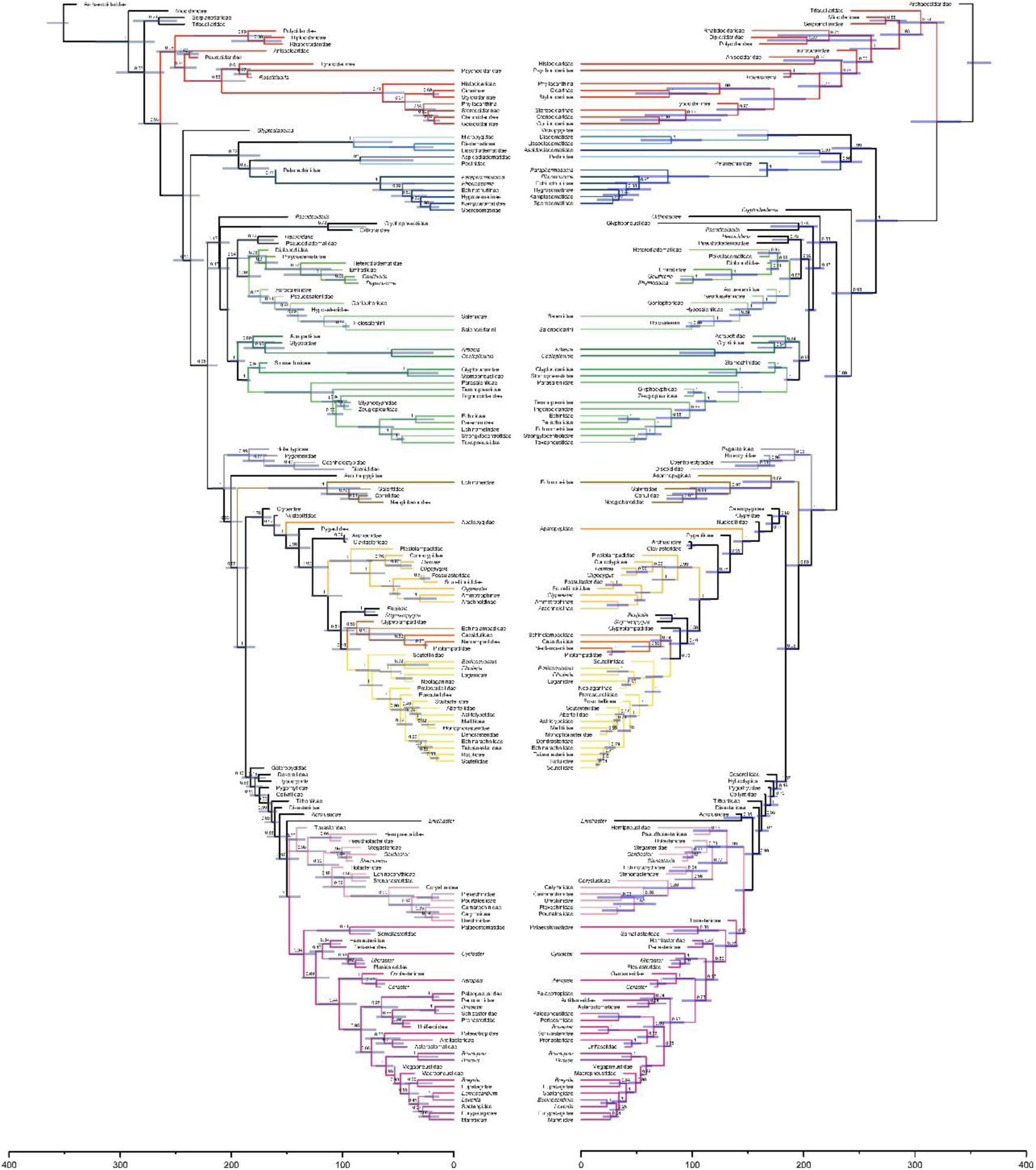
Maximum clade credibility trees for the IGR-SA (left) and TK02-SA (right) inference conditions. The holothurian outgroup was pruned. Branch colors correspond to order-level clades identified in Figure 3A. Blue bars are 95% highest posterior density intervals, numbers on nodes are posterior probabilities. X-axes represent time in millions of years. Terminal branches of length zero for the tree on the left are those shown as direct ancestors in Figure 4A. Beyond the differences highlighted in Figure 3B, it is worth pointing out the different positions of Clypeolampadidae, Galeropygidae, and Toxasteridae.

**Figure S4:**
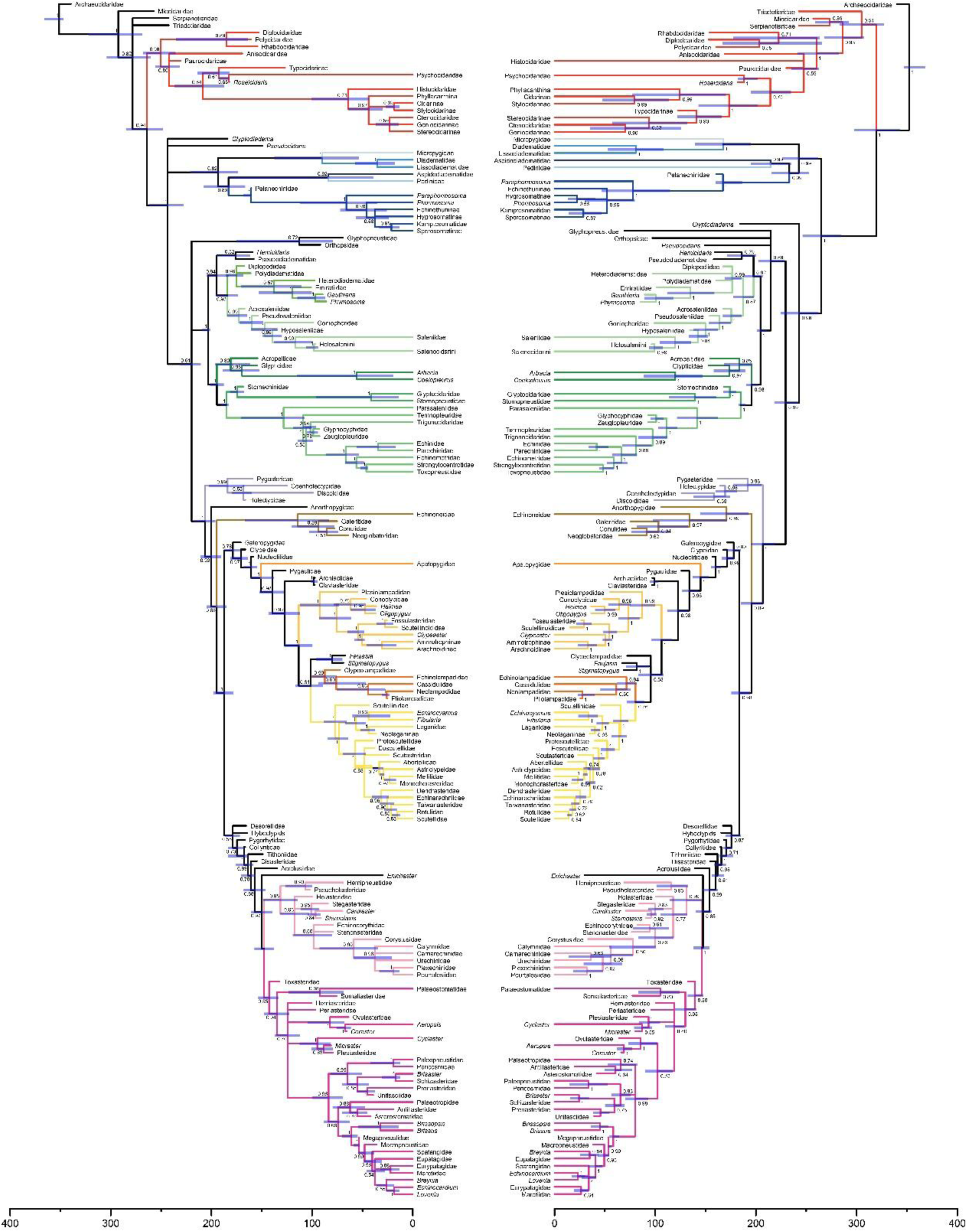
Majority-rule consensus trees for the IGR-SA (left) and TK02-SA (right) inference conditions. The holothurian outgroup was pruned. Branch colors correspond to order-level clades identified in Figure 3A. Blue bars are 95% highest posterior density intervals, numbers on nodes are posterior probabilities. X-axes represent time in millions of years.

**Figure S5:**
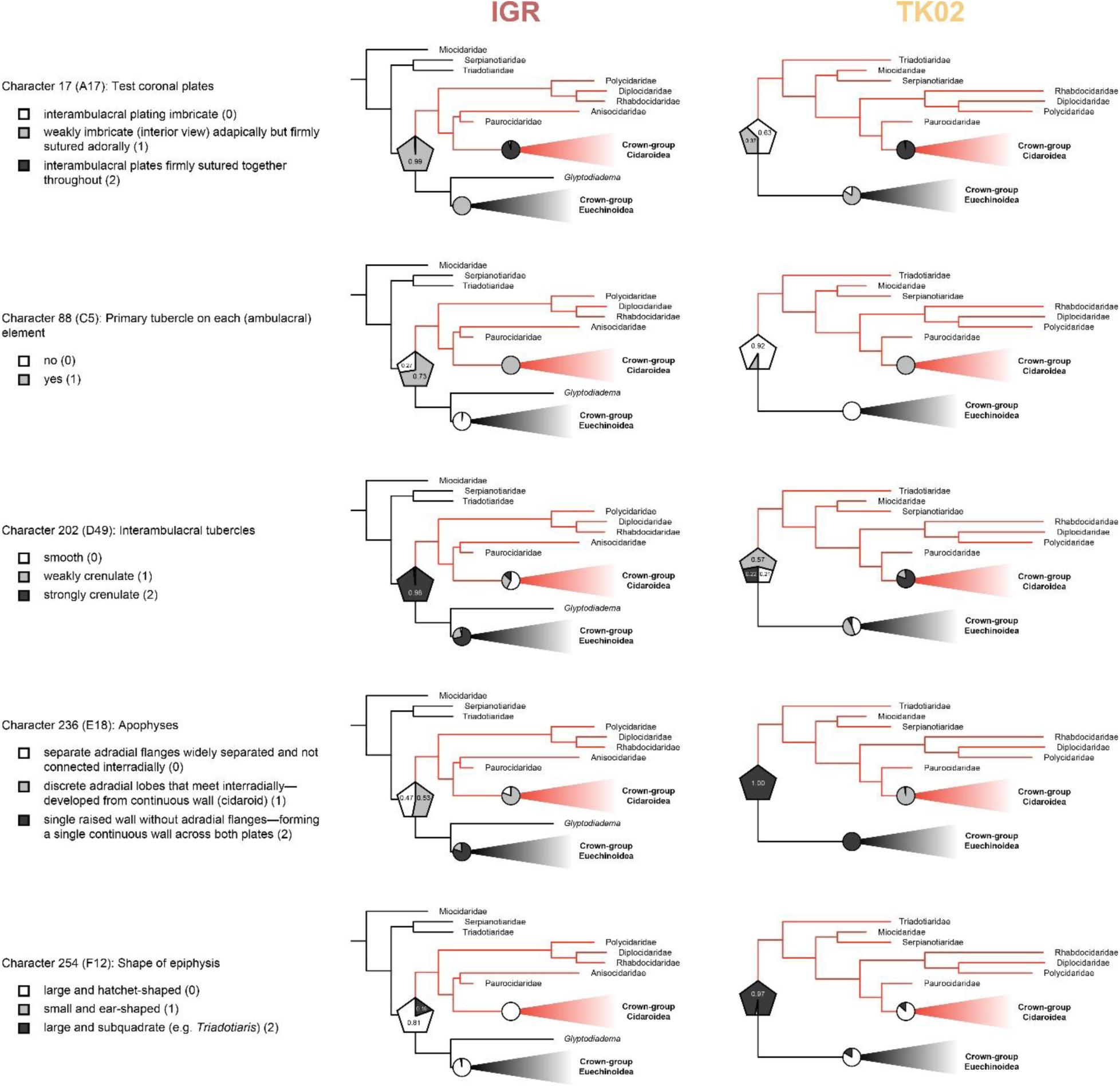
Detailed results of ancestral state reconstructions (ASRs). For thirteen morphological characters, the state reconstructed with the highest likelihood for the node of crown group Echinoidea (depicted with a pentagon) changed between the MCC trees obtained under the IGR-SA (left) and TK02-SA (right) conditions. Five of those characters, highlighted for their relative prominence in the literature, are shown here. Full character and character state descriptions are provided, along with further topological details. ASRs for the nodes of crown group Cidaroidea and Euechinoidea are also shown.

**Figure S6:**
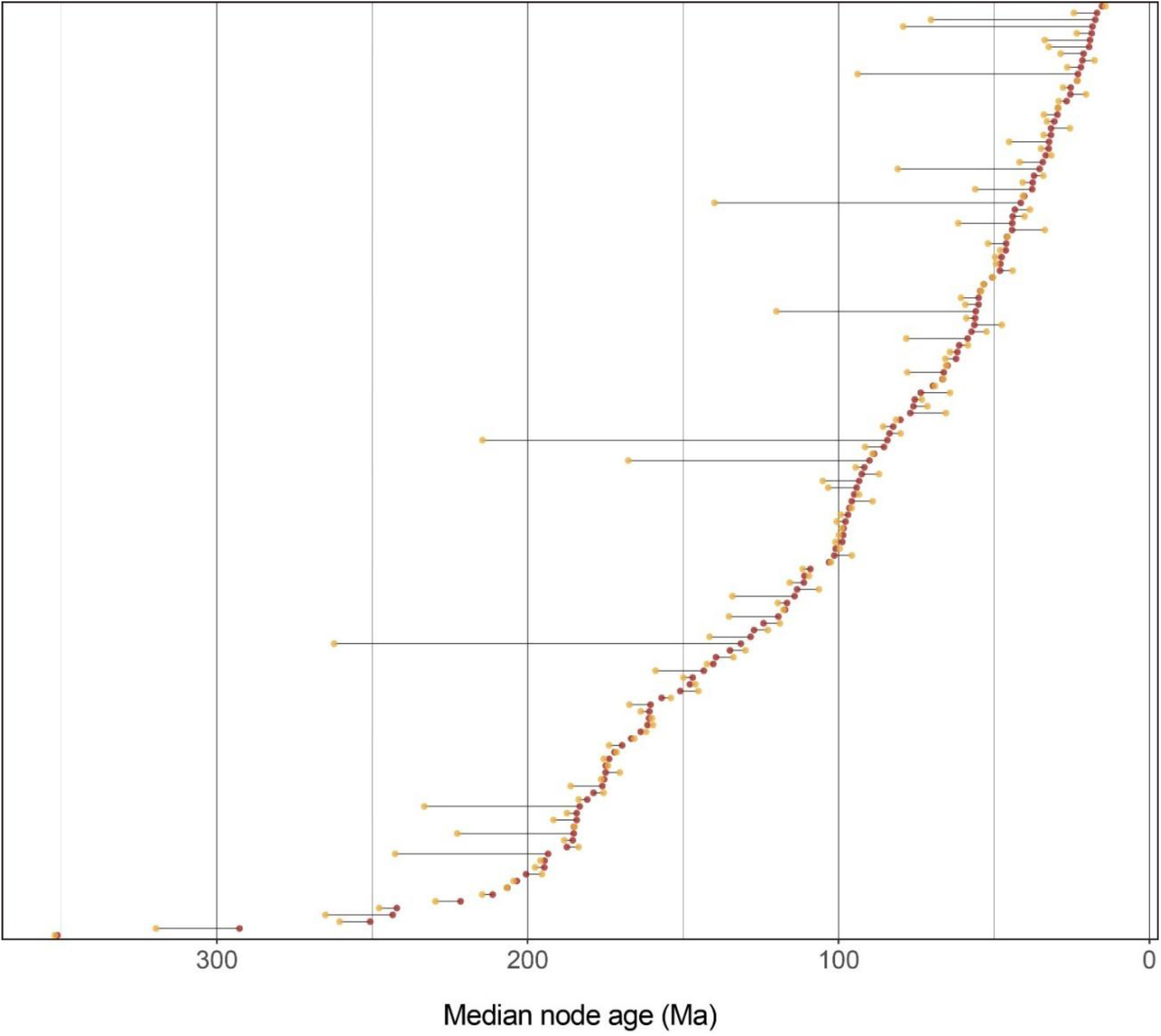
Comparison of median node ages for 138 equivalent nodes (i.e., those giving rise to the same set of tips, regardless of internal topology) in the MCC trees obtained for the IGR-SA (red) and TK02-SA (yellow) analytical conditions (both are shown in Fig. S3).

**Figure S7:**
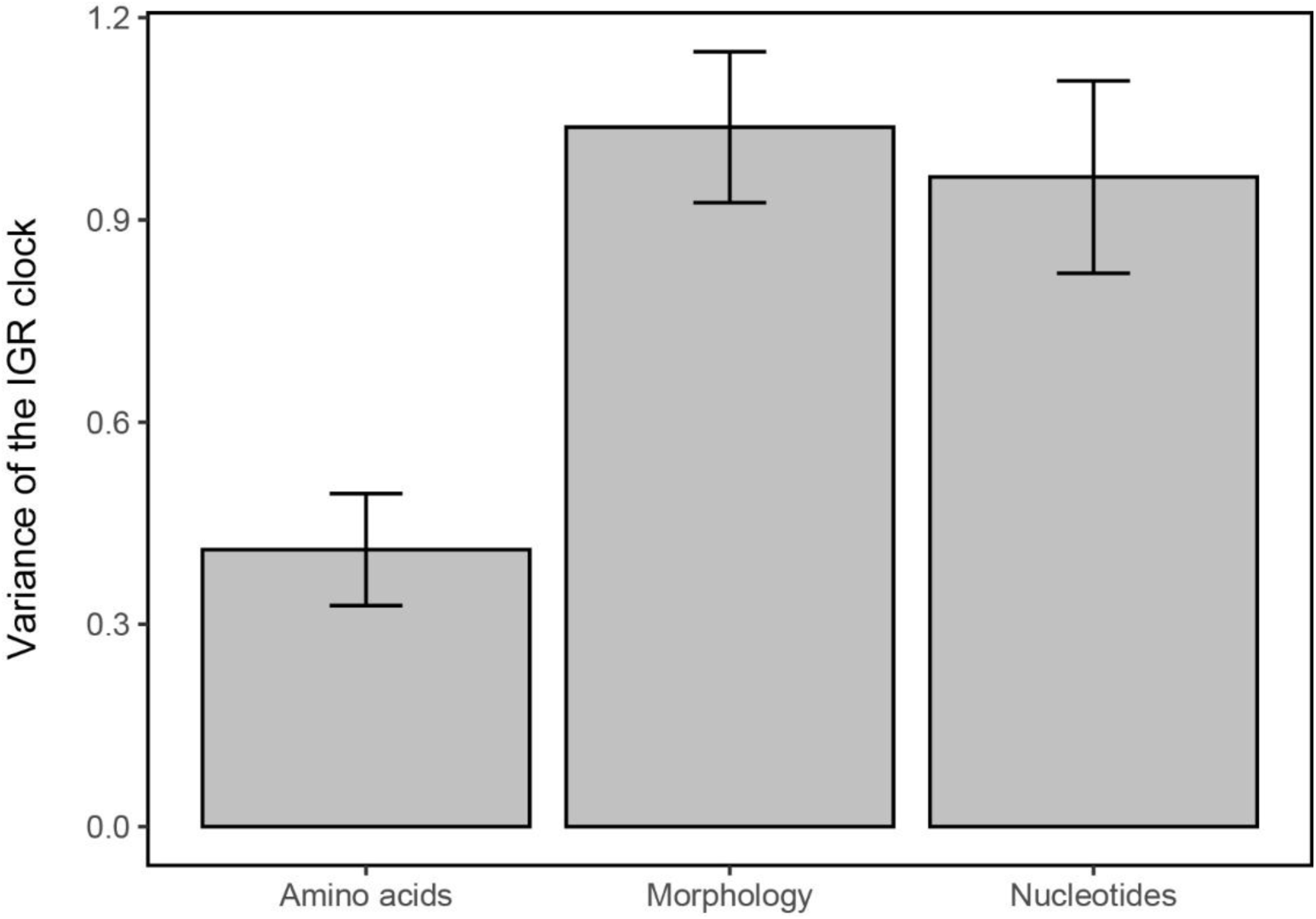
IGR clock variance for each of the three data types. Values shown are posterior means ± one standard deviation. Effective branch lengths are drawn from distributions that are the product of the expected number of substitutions (given the base rate of the clock) and the plotted values. Smaller numbers will result in less branch-specific rate variability. Although only posterior values for the IGR clocks are shown, TK02 variances show the same pattern.

**Table S1:**
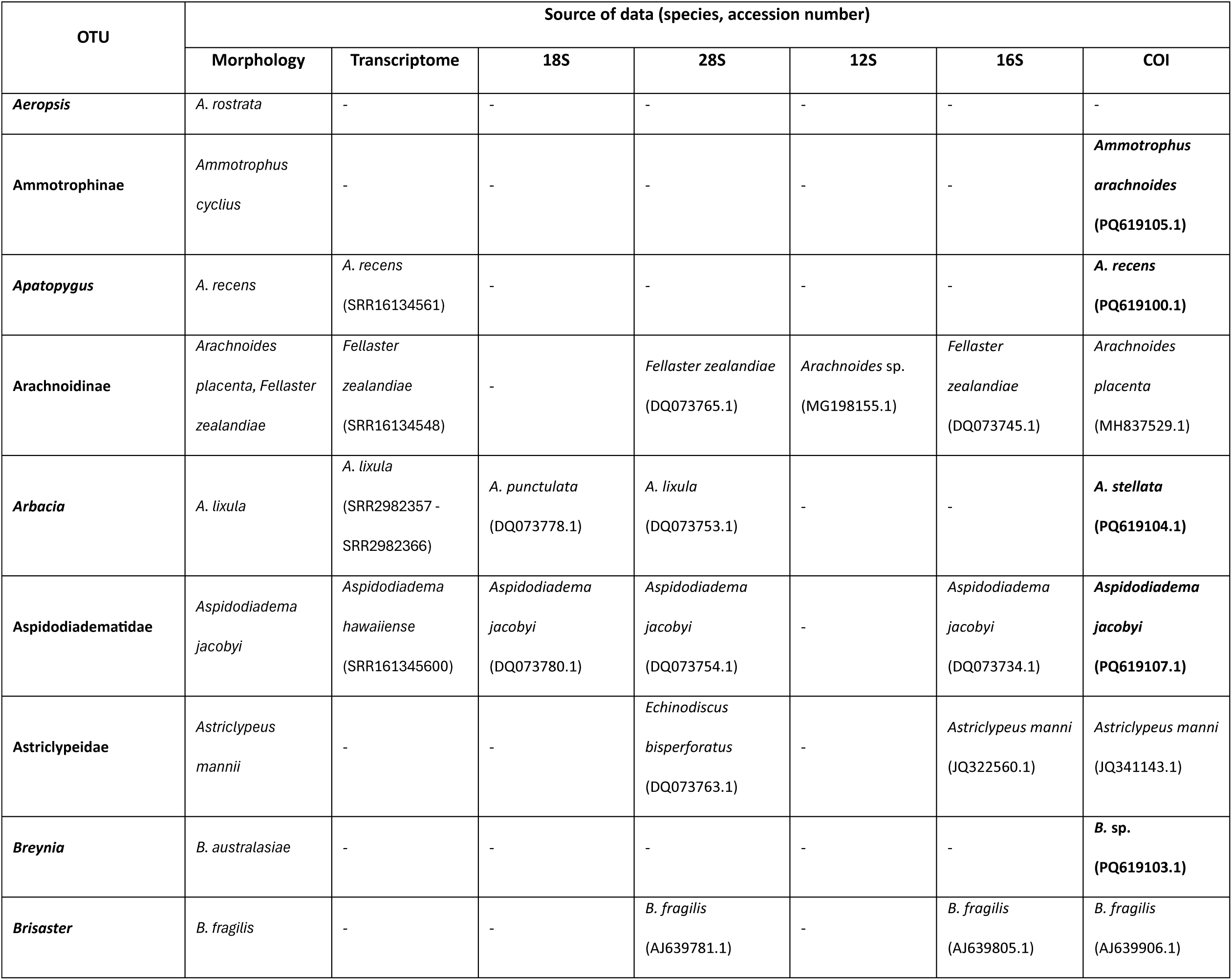
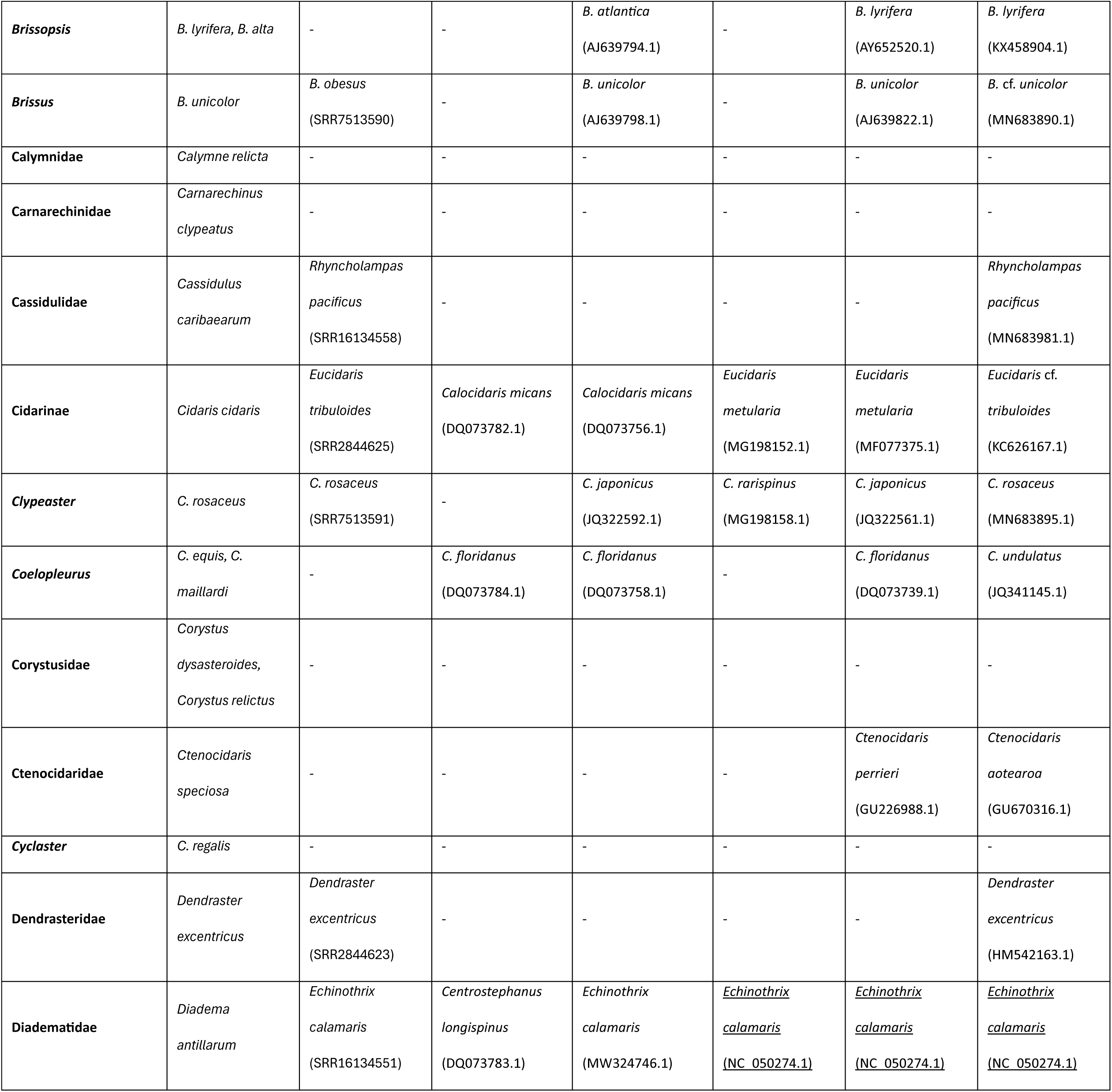
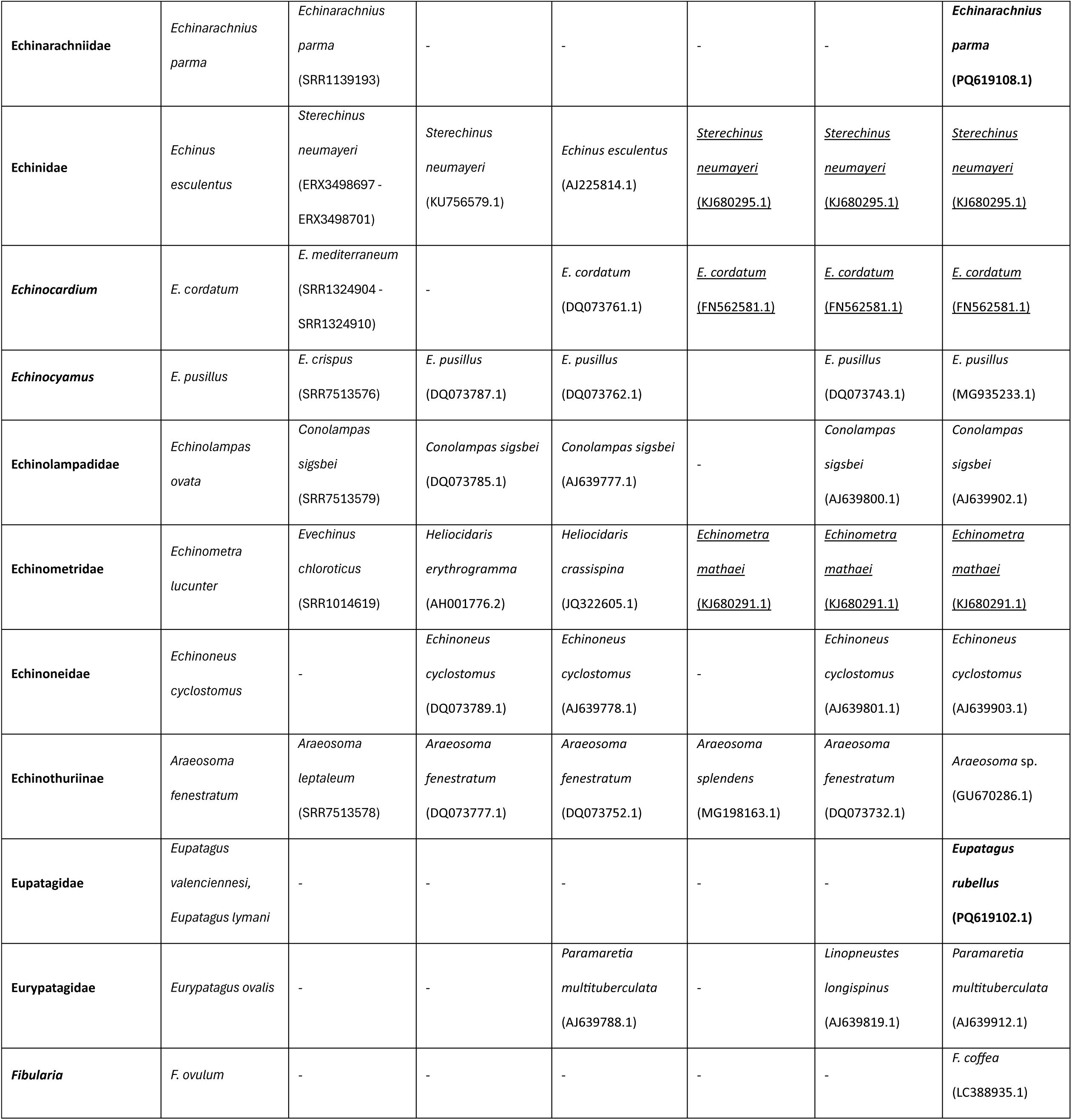
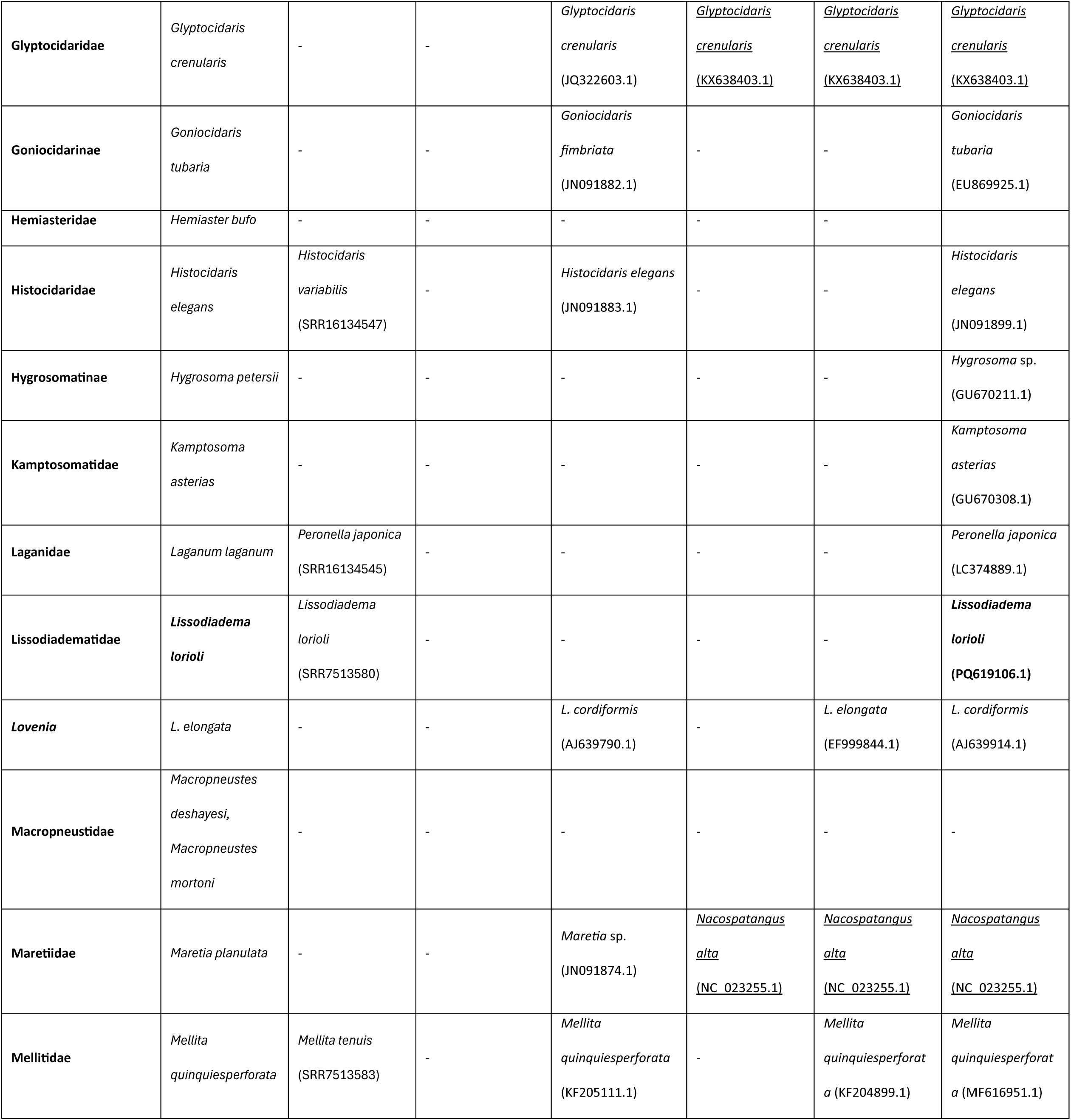
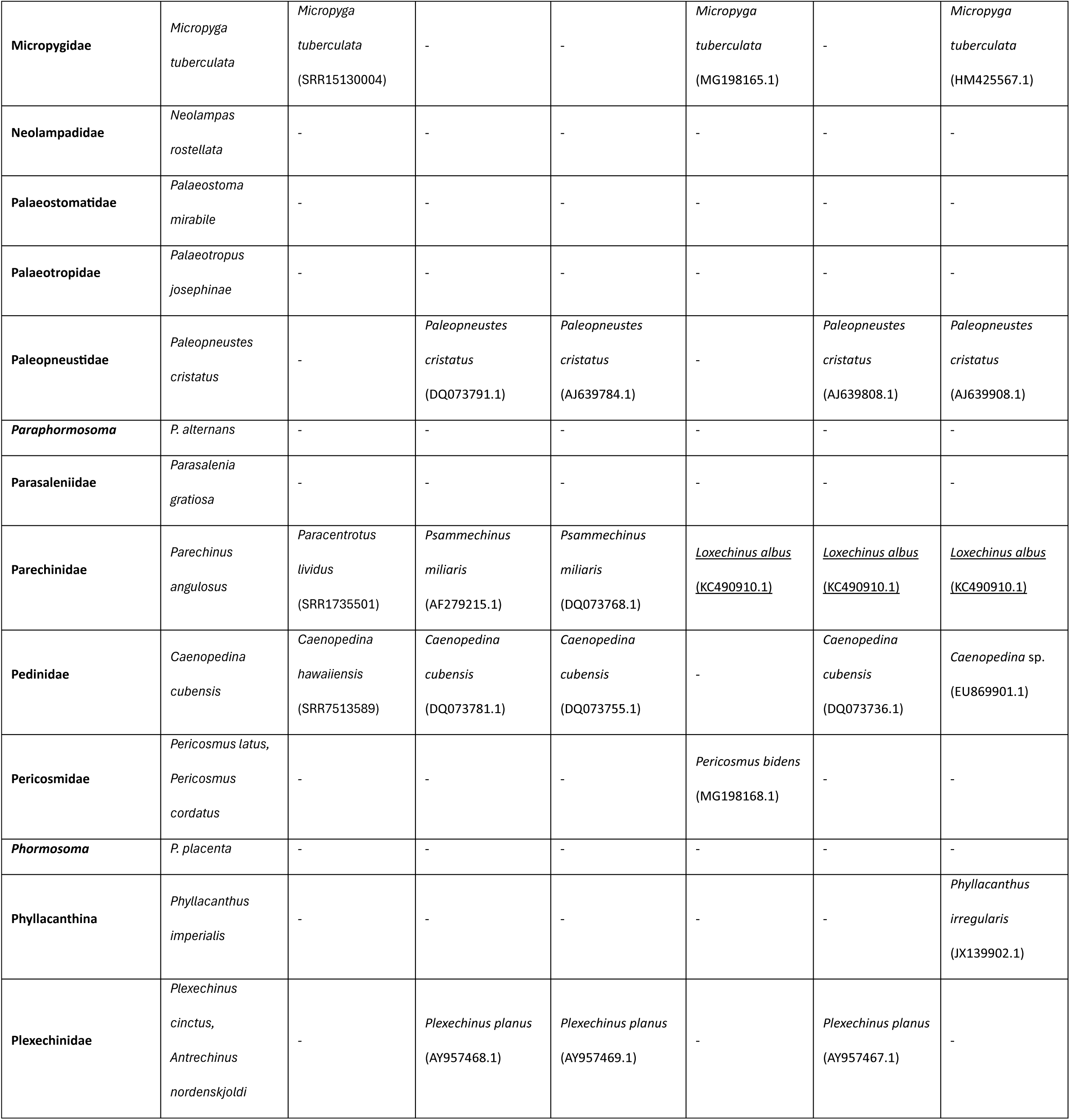
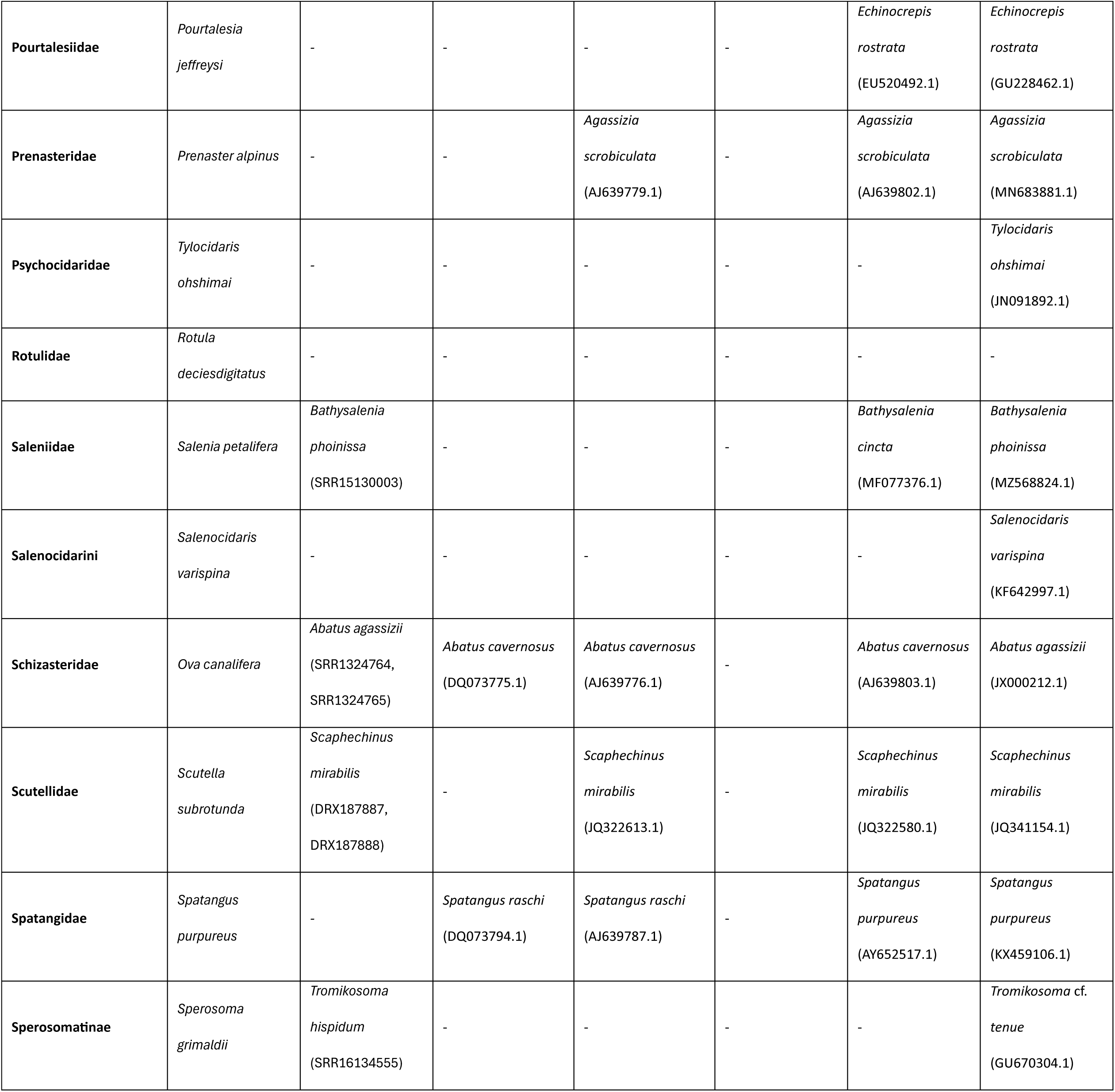
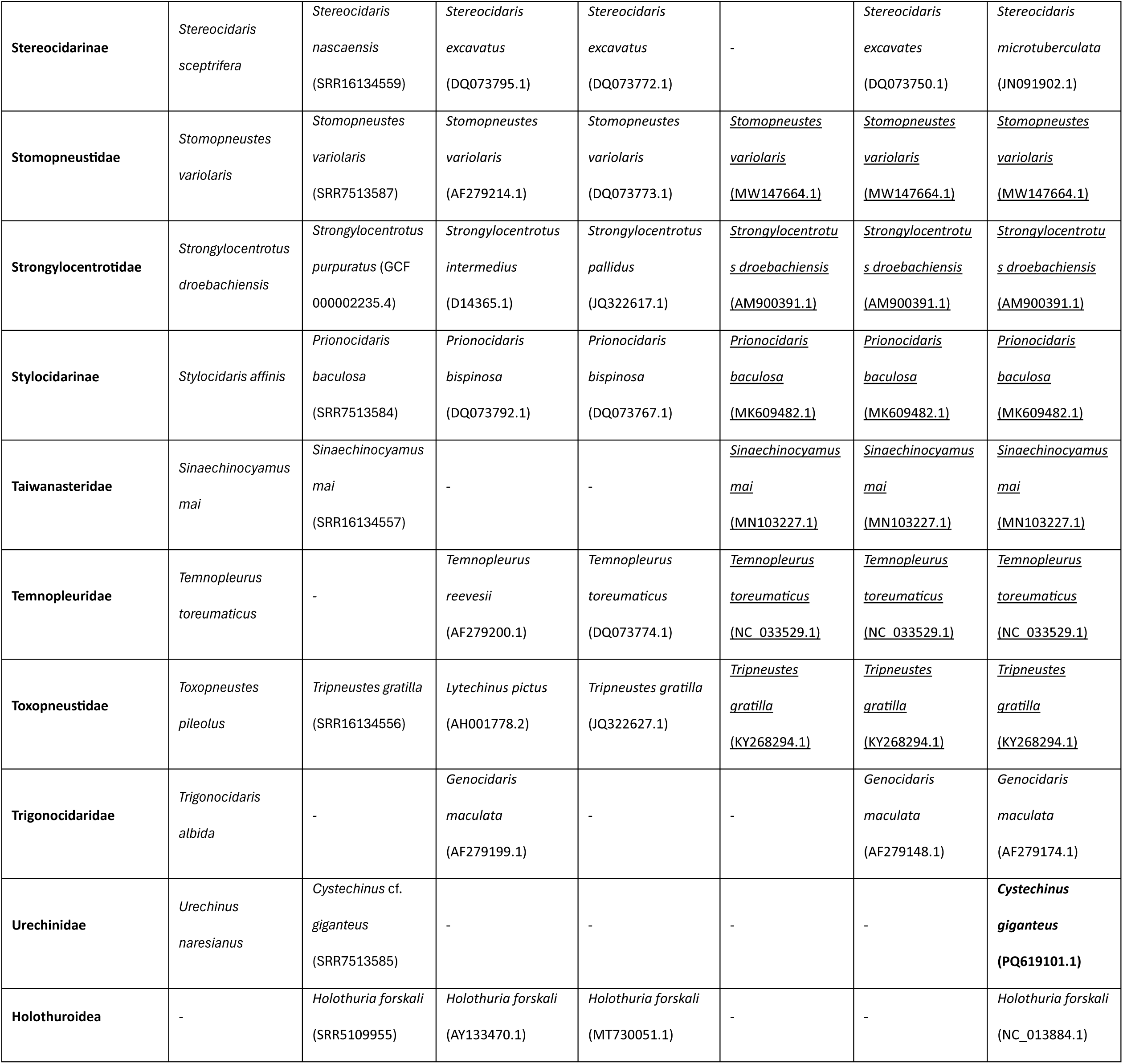
Source of all morphological and molecular data employed. For each operational taxonomic unit (OTU), we list the species used for morphological coding, as well as the source and Genbank/SRA/Genome accession numbers for all molecular data. Entries in bold represent newly generated data, entries in italics represent data extracted from full mitogenomes. Only extant OTUs are listed.

